# Influenza A virus H5Nx genotypes assigned by genetic similarity are incongruent with global phylogenies

**DOI:** 10.64898/2026.10.23.756838

**Authors:** Angela McLaughlin, Hoc Tran, Deanna Tait, Hoang-Hai Nguyen, Josip Rudar, Jolene A. Giacinti, Anthony V. Signore, Oliver A. Lung, Yohannes Berhane, Zvonimir Poljak, Finlay Maguire

**Author notes:** **Corresponding Author:** Finlay Maguire, Faculty of Computer Science Dalhousie University, 6050 University Avenue, PO Box 15000, Halifax, Nova Scotia B3H 4R2.

## Abstract

Influenza A virus H5N1 and related subtypes (H5Nx) have broad host ranges and pose a pandemic risk. Since early 2022, H5 clade 2.3.4.4b has predominated globally, causing unprecedented mortality in wild birds, poultry, and mammals, with spillovers to humans. Accurate viral classification is essential for tracking H5Nx variant emergence and spread, but reassortment among genome segments with distinct evolutionary histories complicates genotyping. H5Nx genotype assignment tools, GenoFlu in North America and Genin2 in Europe, assign segments to clusters by genetic similarity to reference sequences and define genotypes as constellations of segment clusters. However, genetic similarity does not necessarily imply recent shared ancestry due to independently re-occurring mutations and variability in the evolutionary rate. Furthermore, similarity is defined relative to references that may inadequately represent diversity. We evaluated the congruence of genetic similarity-based H5Nx clusters and genotypes with phylogenies of global sequences across eight segments. H5Nx genomes from 41,366 samples were assigned GenoFlu and Genin2 genotypes, and maximum likelihood phylogenies were inferred for each segment. CD-HIT and phylogenetic TreeCluster clusters were also generated for a range of distance thresholds. Clustering methods were evaluated based on phylogenetic purity, monophyly, and diversity. Reassortment within predominant GenoFlu and Genin2 genotypes was evaluated based on their phylogenetic cluster spans. Expectedly, GenoFlu was largely unable to assign genomes from outside of North America, South America, and Antarctica from 2022 to 2025, whereas Genin2 assignments struggled outside of Europe, Africa, and Western Asia. Only 14.2% of GenoFlu clusters were monophyletic, while Genin2 clusters were more variable across segments (overall 49.3% monophyletic). Despite CD-HIT yielding genetic distance clusters of similar size to TreeCluster, they were less monophyletic (44%) and encompassed greater phylogenetic diversity. We identified several genotypes, such as GenoFlu B3.13, D1.1, and D1.3, with segments that had diversified into separate subclusters. Similarly, GenoFlu genotypes B3.5 and A3, and Genin2 genotypes EA-2020-C, EA-2021-AB, and EA-2022-CH may erroneously combine multiple distinct reassortants. Genetic similarity-based H5Nx genotyping tools are rapid, stable, and sufficient for broad categorization, but may group viruses with distinct evolutionary histories and fail to assign divergent viruses, risking biased surveillance, modeling, and risk assessment.

## Introduction

Since influenza A virus (IAV) H5 clade 2.3.4.4b was introduced to North America in late 2021 and 2022, it has caused widespread infection and mortality in wild birds (Caliendo et al., 2022; Rijks et al., 2022; Avery-Gomm et al., 2024; Giacinti et al., 2024; Lane et al., 2024; McPhail et al., 2024), extensive outbreaks in domestic poultry (Youk et al., 2023; USDA and APHIS 2026), and spillover into diverse mammalian hosts, including marine mammals (Falchieri et al., 2026; Banyard et al., 2024; Uhart et al., 2024), terrestrial mammals (Alkie et al., 2023; Elsmo et al., 2023), dairy cattle (Caserta et al., 2024; Nguyen et al., 2025), and humans (Garg et al., 2024; Jassem et al., 2025). Previous incursions of H5 clade 2.3.4.4 viruses into North America in 2014–2015 were detected in subtypes H5N8, H5N2, and H5N1 (Jhung and Nelson 2015; Murti et al., 2015); unlike the current H5 clade 2.3.4.4b panzootic, they were associated with a more limited host range and did not show the same prolonged, multi-year persistence. These contrasting epidemiological patterns, together with extensive reassortment during the current panzootic, have increased the need for approaches to track H5Nx diversity across the whole genome.

The global and long-lasting multi-sectoral impact of the H5N1 panzootic underlines the need for classification systems that reflect evolutionary relationships and transmission chains. Viral lineages, clades, clusters, and genotypes can be used to connect IAV outbreaks across countries and continents (Sánchez-Rodríguez et al., 2026), infer transmission patterns (Youk et al., 2023), evaluate relative fitness (Signore et al., 2025a) and pathogenicity (Alkie et al., 2025), identify reassortants (Wille et al., 2013; Huang et al., 2014; Berhane et al., 2016; Hill et al., 2017; Lycett et al., 2020; Signore et al., 2025b), and structure epidemiological and statistical models (Harrington et al., 2026; Pekar et al., 2026a).

Due to reassortment of its segmented genome, classification of H5N1 and other H5 subtypes (H5Nx) requires consideration of the distinct evolutionary histories of each segment. IAV are classified into subtypes, such as H5N1, based on the combination of their surface glycoproteins, hemagglutinin (HA) and neuraminidase (NA). These subtypes are defined by antigenic properties, including serological cross-reactivity, and are often supported by sequence identity (Poon 2024). HA types, including H5, are further described using phylogenetic lineage- and clade-based nomenclature. For example, H5 lineage A/goose/Guangdong/1/1996 is divided into multiple clades including 2.3.4.4b (Duan et al., 2008; Smith, Donis, and WHO/OIE/FAO Working Group 2015; Bellido-Martín et al., 2025). H5 clade 2.3.4.4b has diversified considerably since its emergence, with the earliest complete sequences available within H5N8 and H5N2 sampled from China in 2010. Subsequent reassortment yielded H5N1 clade 2.3.4.4b in Europe 2020–2021 (King et al., 2022), which rapidly spread across Europe, Asia, Africa, and North America, generating numerous reassortants (Xie et al., 2025).

Several H5Nx genotyping systems have been developed to track the emergence, spread, and reassortment of H5 clade 2.3.4.4b viruses. In North America, GenoFlu assigns segment-level cluster labels using genetic similarity (the proportion of nucleotides shared between query and reference), then genotypes are defined by the constellation of segment-level labels (USDA-VS 2026; Youk et al., 2023). This framework has supported the characterization of multiple incursions of H5 clade 2.3.4.4b from Eurasia to North America via the Atlantic and Pacific, and as well as the identification and tracking of subsequent reassortant genotypes (Youk et al., 2023; Signore et al., 2025a; Harrington et al., 2026).

Similarly, Genin2 was developed to assign H5Nx clade 2.3.4.4b viruses to genotypes based on genetic similarity to European reference sequences (Fusaro et al., 2024). Its assignment algorithm uses machine learning models trained on European genotypes designated on an initial phylogeny. These genotypes have been used for genomic epidemiology and surveillance for example, to describe H5N1 and H5N8 collected from wild bird and poultry in Egypt from 2022–2024 as European Genin2 genotypes EA-2021-AB and EA-2020-A (Eid et al., 2025). A phylogeographic analysis of H5 sequences from Italy identified multiple introductions of EA-2022-BB and EA-2021-AB that contained multiple minor genotypes nested within them on the HA tree, suggesting genotypes did not correspond with distinct outbreaks (Fornasiero et al., 2026). Regional genotype classifications from GenoFlu and Genin2 are also cross-referenced by ggFlu (Song 2026), which maps queries to predefined genotypes by placing sequences by parsimony on segment reference trees.

Both GenoFlu and Genin2 genotyping tools rely on genetic similarity to a set of references to assign segment clusters. Genetic similarity-based assignment offers several practical advantages, including computational efficiency, scalability, and relative stability. Depending on the downstream application, genetic similarity clustering is sufficient for identifying rough, easily communicated, genetic groupings for outbreak response. For example, the initial H5N1 clade 2.3.4.4b spillover into dairy cattle was associated with genotype B3.13, while subsequent D1.1 outbreaks in cattle were genetically distinct and resulted from independent introductions (Caserta et al., 2024; Pekar et al., 2026b). When the references adequately represent population diversity, the most genetically similar reference to a sequence may also be the closest phylogenetic relative; however, this correspondence is not guaranteed. Genetically similar sequences are not always evolutionarily proximate due to convergent evolution or homoplasy, where the same mutation occurs multiple times across the tree (Wagner 2014; Guo et al., 2019). Variable evolutionary rates (Fleming, Valero-Gracia, and Struck 2023), incomplete or biased sampling (Bush et al., 2000), and reference selection can further weaken the correspondence between genetic similarity and evolutionary relatedness. Alignment uncertainty and the treatment of gaps and insertions/deletions can also affect estimates of dissimilarity and phylogenetic relationships (Rivas and Eddy 2008). As a result, segment labels designated or assigned by genetic similarity may not always correspond to monophyletic clades, in which all tips descending from the cluster’s most recent common ancestor (MRCA) carry the label; genotypes built from those labels may combine viruses that are genetically but not phylogenetically close. By contrast, phylogenetic cluster designation and assignment are based upon sequences’ shared ancestry depicted as phylogenetic trees, explicitly incorporating rate variation and convergent evolution. This approach also enables identifying reassortants based upon discordant evolutionary histories across segments (Markin et al., 2025). Despite these strengths, phylogenetic clustering is computationally demanding, carries uncertainty, and poses challenges for cluster stability as new genomes become available. An ideal H5Nx genotyping approach should describe global whole genome diversity, remain adaptable to emerging diversity, and define segment clusters using clear and reproducible criteria reflecting evolutionary history.

Our objective was to evaluate the extent to which H5Nx segment cluster and genotype assignments generated by tools based on genetic similarity correspond to phylogenetic trees across the global diversity of all eight genome segments. We compared assignments from GenoFlu and Genin2 with phylogeny-based clustering using TreeCluster (Balaban et al., 2019) and genetic similarity clustering using CD-HIT (Li and Godzik 2006) across a range of thresholds for a global H5Nx dataset. These analyses provide a framework for evaluating the extent to which H5Nx genotype labels capture recent shared ancestry across the genome, as well as for distinguishing reassortment from mutational drift. Segment cluster constellations based upon shared recent ancestry provide a more explicit evolutionary basis for interpretation of genotypes and a clearer framework for characterizing reassortment. Evolutionarily accurate genotypes are critical for influenza surveillance and response.

## Methods

### Data retrieval and cleaning

On December 10, 2025, nucleotide sequences and metadata were downloaded from the National Center for Biotechnology Information (NCBI) Virus database (Bao et al., 2007) for all available IAV H5Nx sequences, excluding lab passaged or vaccine strains, collected since January 1, 1961 (n=19,449 samples, n=174,051 nucleotide sequences). Data were also downloaded from the Global Initiative on Sharing Avian Influenza Data (GISAID) EpiFlu (Shu and McCauley 2017) database on December 10, 2025, including sequences from type A influenza, filtering to H5 only, from all hosts and locations, with all eight segments, collected from the earliest sample (January 1, 1975) to the day of data download (n=42,415 samples, n=363,401 nucleotide sequences).

Samples were excluded if they were under a publication embargo for more than three months on the day of data download, or if the strain name included ‘vaccine’, ‘passaged’, ‘mutant’, ‘lab derived’ or ‘clone’ (Fig. 1). GenBank title and isolate name were parsed to fill in and cross-check missing, misspelled, and mislabeled metadata including sampling location, host, collection date, and segment. Samples were excluded if they were missing host after parsing. Segment sequences were excluded if they were duplicated within a sample (the sequence with higher coverage or more recent date was kept) or if a sample did not have all eight segment sequences.

**Figure 1.**
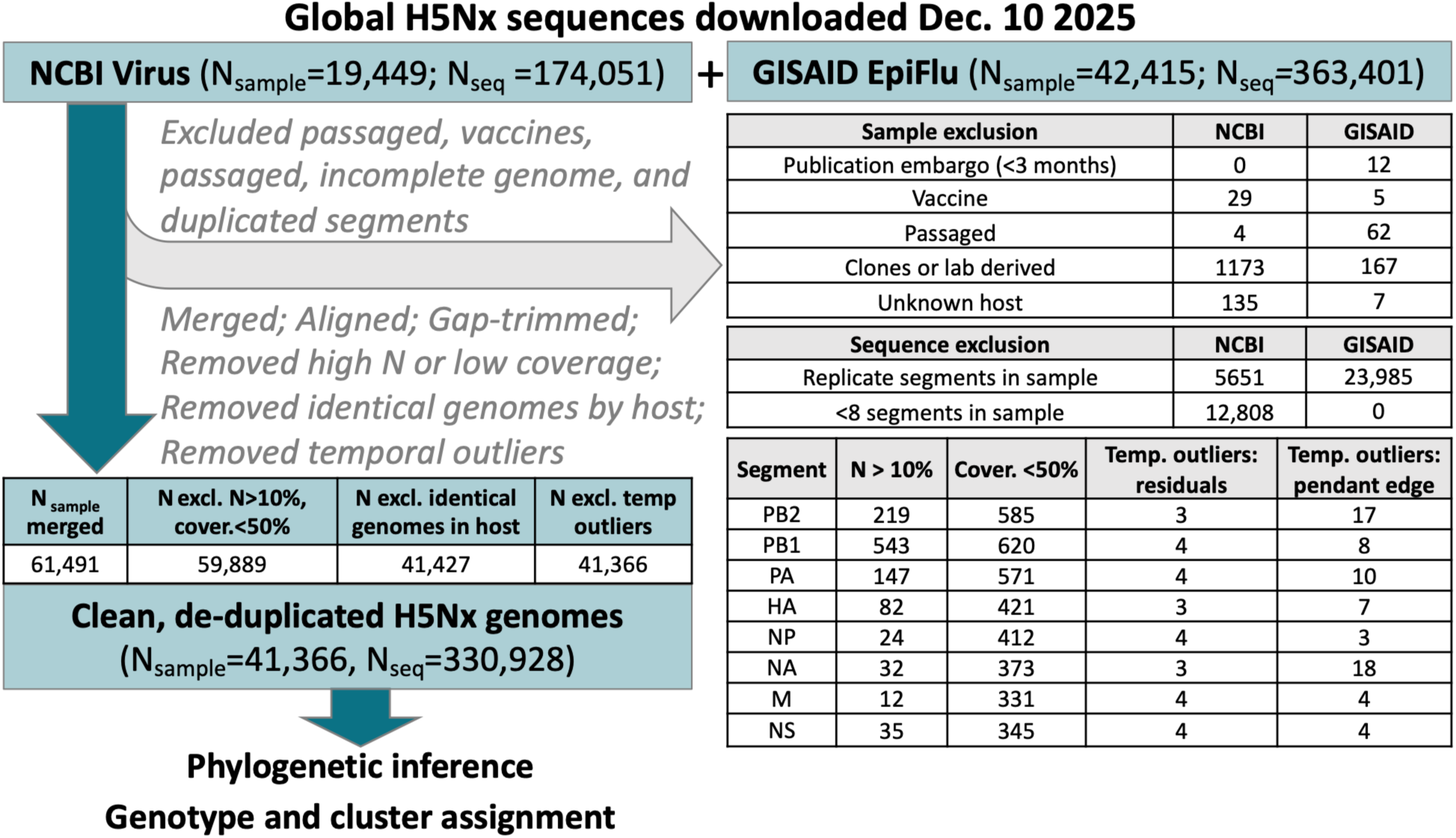
Data cleaning pipeline for global H5Nx genetic sequences from GISAID EpiFlu and NCBI Virus. Nucleotide sequences and metadata were cleaned to exclude samples if they were under publication embargo, vaccine strains, passaged, clones, lab-derived cultures, or had unknown host. Segment sequences were excluded if there were duplicate segments within a sample or fewer than eight segments. Data were merged, aligned, and gap trimmed. Segment sequences were excluded if ambiguous (N) sites exceeded 10%, or coverage was lower than 50% of trimmed width. Data were de-duplicated of identical whole genome sequences within common hosts. Preliminary phylogenetic trees for each segment were used to identify and prune temporal outliers with high residuals or mutations on the pendant edges. Samples were excluded if any of their segment sequences were excluded. The cleaned and de-duplicated H5Nx samples and sequences were utilized to infer maximum likelihood phylogenies for each segment and assign genotypes and clusters.

Merged NCBI and GISAID nucleotide sequences were aligned using MAFFT version 7.490 (Katoh and Standley 2013). Alignments were trimmed using the R package ‘Biostrings’ version 2.78.0 (Pagès et al., 2020). First, a contiguous region bounded by positions with at least 10% non-gap sequences was identified, and all positions outside this region were removed to trim frayed ends; then, internal positions were trimmed if more than 99% of sequences had a gap, retaining insertions and deletions with at least 1% frequency. Final trimmed widths were reported (Table S1). Sequences in trimmed alignments were excluded if they had >10% ambiguous sites or <50% coverage (Fig. 1), to balance the needs for high-quality sequences with maximal inclusivity. Samples were excluded if any of their segment sequences were excluded. Identical whole genome sequences originating from the same host were identified as duplicates; within each duplicate group, sample identifiers were sorted alphabetically, and the first identifier was retained.

Sampling locations were converted to global geographic regions using R package ‘countrycode’ version 1.6.1 (Arel-Bundock, Enevoldsen, and Yetman 2018) based on the United Nations M49 ‘geoscheme’ (Table S4). Note that within this scheme, Mexico is grouped with Central America, Kazakhstan with Central Asia, and Greenland with Northern America. Global maps were generated using R packages ‘sf’ version 1.0.24 (Pebesma 2018), ‘rnaturalearth’ version 1.2.0 (Massicotte and South 2026), and ‘ggplot2’ version 4.0.2 (Wickham 2016). Maps use a Robinson projection (central meridian 0° with 20° graticule).

### Maximum likelihood phylogenetic inference

Trimmed nucleotide alignments were used to infer preliminary approximate maximum likelihood trees using FastTree version 2.2.0 (Price, Dehal, and Arkin 2010) with a generalized time-reversible (GTR) model, rooted with root-to-tip (RTT) linear regression in R package ‘ape’ version 5.8.1 (Paradis and Schliep 2018). Sequences were pruned as temporal outliers if their RTT residuals exceeded a maximum residual of 10 standard deviations, or if their pendant edge (terminal branch preceding tip) length exceeded a maximum number of mutations, set to 6% of segments’ trimmed nucleotide alignment widths (Table S1). Adjusted R-squared values for the RTT regression were compared before and after excluding temporal outliers (shown for HA, Fig. S1). Pruned starting trees and alignments were used as input for IQ-TREE version 2.0.4 (Minh et al., 2020) with a GTR+F+R10 substitution model, as supported by ModelFinder (Kalyaanamoorthy et al., 2017) for most segments and rooted using RTT regression. Unassigned H5 clades (Fig. S4) and NA subtypes (Fig. S5) were imputed based on the phylogenetic neighborhood: as the clade label defined by its parental node if it defined a monophyletic clade, or as the label of its nearest neighbor.

### Cluster and genotype assignment and phylogenetic evaluation

Sequences were assigned by genetic similarity to genotypes using GenoFlu version 1.06 (Youk et al., 2023; USDA-VS 2026) and Genin2 version 2.1.6 (Fusaro et al., 2024; Sartori 2026), and segment cluster labels were parsed from genotype constellations where available. For instance, GenoFlu genotype B3.13 is the alias for constellation, ‘PB2-am2.2_PB1-am4_PA-ea1_HA-ea1_NP-am8_NA-ea1_M-ea1_NS-am1.1’, which includes PB2 cluster am2.2.

Sequences were assigned to phylogenetic clusters in segment trees using TreeCluster v1.0.4 ‘max-clade’ (Balaban et al., 2019), which identifies sequences within a clade connected by pairwise patristic distances less than or equal to a threshold, for which we evaluated from 0.01 to 0.25 substitutions/site (subs/site). The optimal patristic distance threshold for this dataset was identified based on yielding a stable number of clusters and singletons (i.e., at the elbow of the curve, Fig. S6) and a maximum cluster size less than ∼25% of the dataset size, a boundary chosen based on the observation that higher cut-offs yielded clusters with >10,000 sequences. TreeCluster cluster identifiers (IDs) were numbered from bottom to top of phylogenies ladderized in R package ‘ggtree’ version 4.0.4 (Yu et al., 2016). Singletons were not distinguished within genotype constellations nor counted towards clusters. To control for differences in the number and size of clusters in comparing genetic and phylogenetic clustering, we identified CD-HIT clusters (Li and Godzik 2006) connected by 85–99.5% genetic similarity utilizing a similar approach as above to identify an optimal threshold based on the distribution of cluster sizes.

We evaluated the diversity and composition of predominant GenoFlu and Genin2 genotypes relative to TreeCluster (Fig. 5). We compared labels on divergent segment subtrees pruned at the MRCA of GenoFlu genotypes D1.1 (Fig. S11) and B3.13 (Fig. S12), and Genin2 genotypes EA-2020-C (Fig. S13) and EA-2022-BB (Fig. S14). Within major genotypes, we report the number of comprising phylogenetic clusters, their cluster ID range (minimum to maximum ID), cluster ID span (maximum ID – minimum ID + 1), and the span-to-cluster ratio (cluster ID span/number of clusters; Table S5, Table S6). Whereas a span-to-cluster ratio near 1 indicates continuous mutational drift, a higher ratio (>2) may suggest the presence of multiple reassortants erroneously grouped together because clusters are discontinuous (Fig. S22). Singletons were excluded from the cluster count, span, and ratio.

Across clustering methods, we compared the number of clusters, median cluster size, maximum cluster size, median cluster purity (proportion of tips with cluster label descending from the MRCA of all tips with cluster label, excluding unassigned), proportion of monophyletic clusters (with 100% purity), and median phylogenetic diversity (PD; Faith 1992), calculated as the sum of branch lengths (substitutions/site) for the minimum spanning subtree connecting clusters’ sequences. To account for differences in cluster size, we also compared the proportion of monophyletic clusters (Fig. S18), median purity (Fig. S19), and median PD (Fig. S20) across methods within cluster size bins (2, 3–5, 6–10, 11–50, 51–200, and >200 sequences).

## Results

### Summary of cleaned H5Nx genomes and metadata

After H5Nx sequences were cleaned, filtered to whole genomes, de-duplicated of identical genomes within hosts, and pruned of temporal outliers, there were 330,928 sequences across eight segments from 41,366 samples in the dataset (Fig. 2). Collection dates ranged from 1961 (H5N3 from a tern in South Africa; NCBI Virus) to November 30, 2025, though we focus on sequences since 1996.

**Figure 2.**
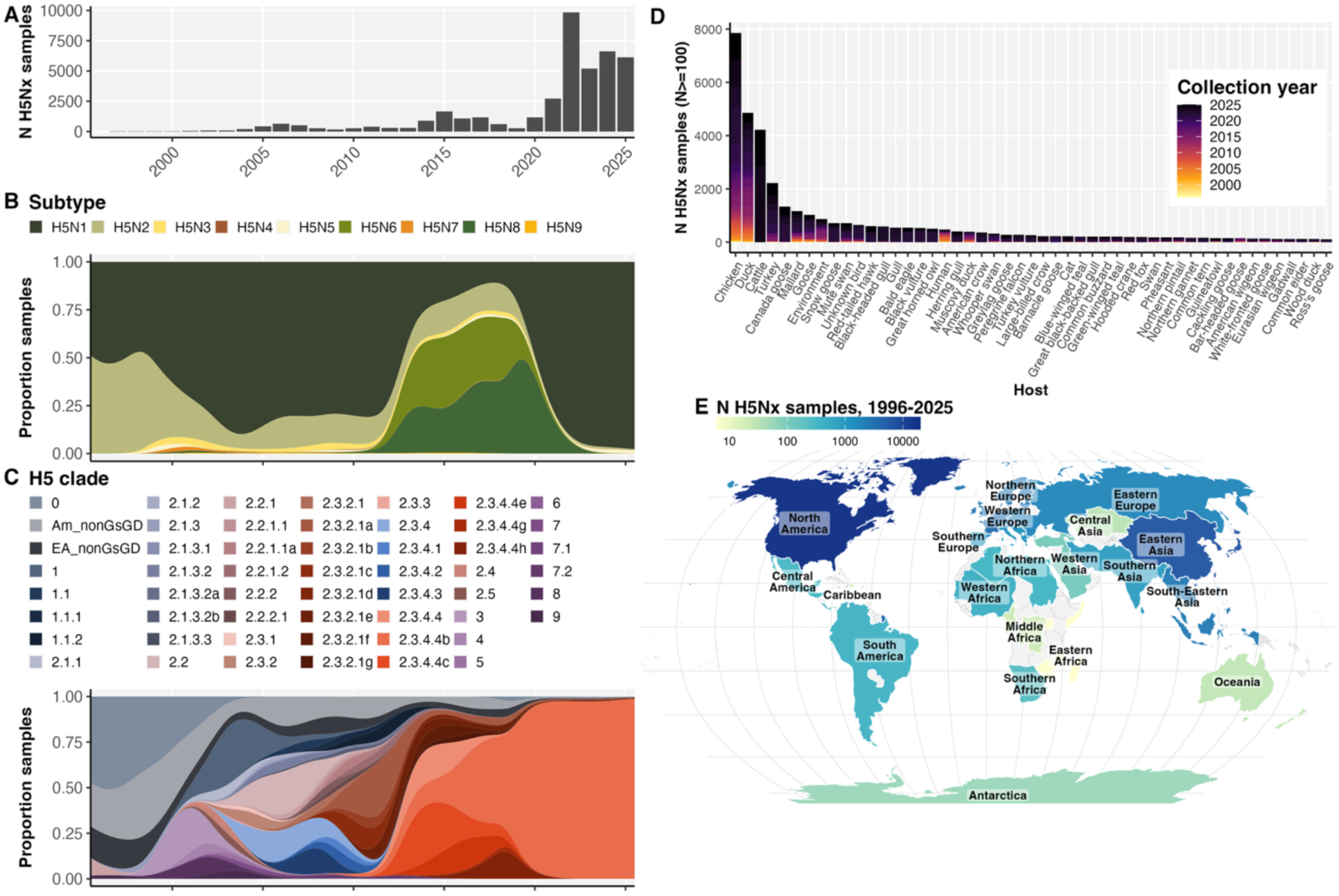
Cleaned and de-duplicated H5Nx samples with whole genome sequences from 1996–2025. Distributions of A) clean H5Nx samples across collection years; proportion of samples by B) subtype and C) H5 clade; D) frequency of hosts (lowest taxonomic level, Table S3) with at least 100 samples since 1996; and E) samples available from global regions since 1996. Equivalent plots with full dataset since 1961 (Fig. S2).

H5N1 has been the most common H5Nx subtype globally since 2022, having previously predominated from 1999 to 2013 until being displaced by H5N8 and H5N6 between 2014 and 2021 (Fig. 2B). H5 clade 2.3.4.4b has been the predominant clade (after removing identical sequences within each host) since 2016, and the majority clade since 2020 (Fig. 2C). Labeling NA types and H5 clades on phylogenies revealed sequences with incongruently labeled NA types and H5 clades (Fig. S4 and Fig. S5). For example, there were ‘EA_nonGsGD’ labels nested within H5 clade 2.3.4.4b (Fig. S4), N8 and N5 types nested in N1, and N8 nested in the N6 clade (Fig. S5). Sequences were collected across all global regions, except Melanesia, though mostly from North America in 2022–2025, followed by Eastern Asia, South-Eastern Asia, and European regions since 2020 (Fig. 2E). Chicken, duck, cattle, turkey, Canada goose, and mallard were the most frequently sampled hosts (Fig. 2D). Cattle predominated prior to removing duplicate genomes by host. After accounting for spelling errors and redundancy, and filtering out environment, pet food, and dairy products, there were 537 unique hosts, including 442 species, in the H5Nx dataset (Fig. S3; Table S3). Within samples designated as H5 clade 2.3.4.4b (after filling in missing clade labels), there were 464 hosts and 384 species, more than previously appreciated.

### Global assignment gaps in H5Nx genotyping systems

Neither GenoFlu nor Genin2 genotypes were assigned at high rates across H5Nx sequences from all global regions from 2020–2025 (Fig. 3). There were 31,683 sequences collected in 2020–2025, of which 63.8% were assigned GenoFlu genotypes, while 34.4% were assigned Genin2 genotypes. As expected, the GenoFlu assignment rate was highest in North America, and high in the Caribbean, South America, and Antarctica, corroborating these regions’ ecological and epidemiological connections. The Genin2 genotype assignment rate was expectedly highest in European regions, and high in Western and Middle Africa (though lower in Northern Africa), and Western, Central, and Eastern Asia. Sequences from Southern and South-Eastern Asia, Central America, and Oceania had low assignment from both tools.

**Figure 3.**
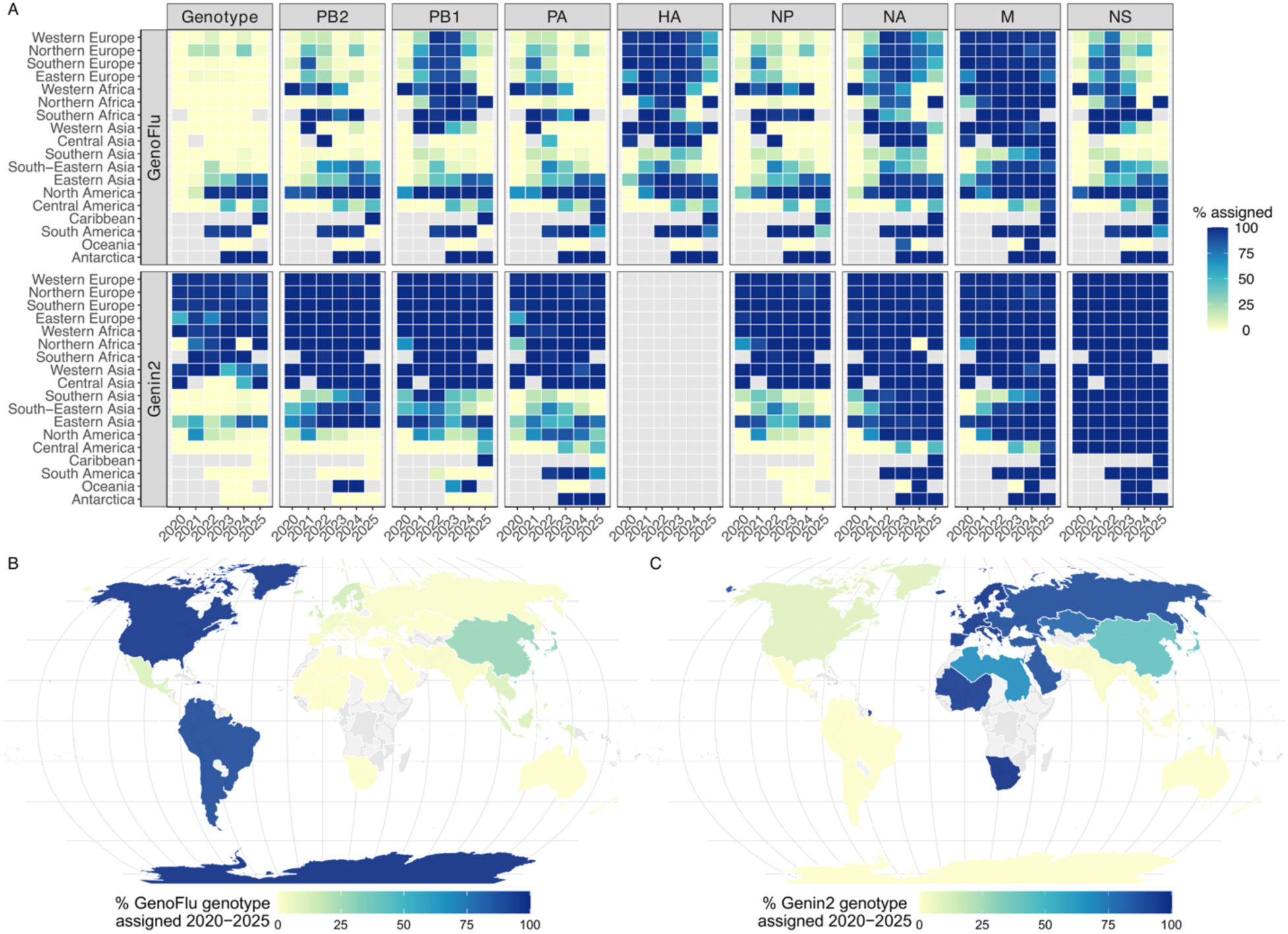
Genotype assignment by global region for GenoFlu and Genin2 in 2020–2025. A) Heatmaps depict percent assignment for genotypes and segment clusters for each region and year. Grey cells indicate no data. Genin2 does not distinguish HA clusters. UN subregions shaded by the percentage of sequences assigned genotypes from 2020– 2025 for B) GenoFlu, and C) Genin2. Polygons are Natural Earth country boundaries dissolved to UN subregions; white lines mark subregion borders, and only countries with sequences in 2020–2025 are shaded. Grey countries/regions indicate no sequences in 2020–2025. Maps depict a Robinson projection (central meridian 0° with 20° graticule).

Incomplete genotype assignment was driven by unevenly assigned segment clusters. For example, North American sequences had lowest Genin2 assignment in PB2, PB1, and NP, but NS was fully assigned, and NA and M were mostly assigned. European sequences’ GenoFlu assignment was high for HA and M, but low for PB2, PA, and NP, and moderate for PB1, NA, and NS (highest in 2022 and 2023).

### H5Nx clusters assigned using genetic similarity are inconsistent with shared recent ancestry

Next, we considered to what extent genotypes and clusters assigned using genetic similarity shared recent ancestry. We detected 86 GenoFlu genotypes, including 33 major and 53 minor genotypes. Across segments, GenoFlu assigned relatively few clusters (6–34 clusters per segment), spanning a wide range of sizes from 2–21,707 tips, with 52% of sequences in a single HA cluster. GenoFlu cluster labels were non-monophyletic across all eight segment trees (Fig. 4, Fig. S10). For example, on the HA tree, ea4 was polyphyletic, covering 5 disconnected clades. Additionally, GenoFlu clusters had poor separation of within- and between-cluster patristic distances (Fig. S15); in PB2 there were within-cluster patristic distances of 0.3 subs/site, which equates to 702 nucleotide substitutions within a cluster (PB2 trimmed width: 2341 nucleotides).

**Figure 4.**
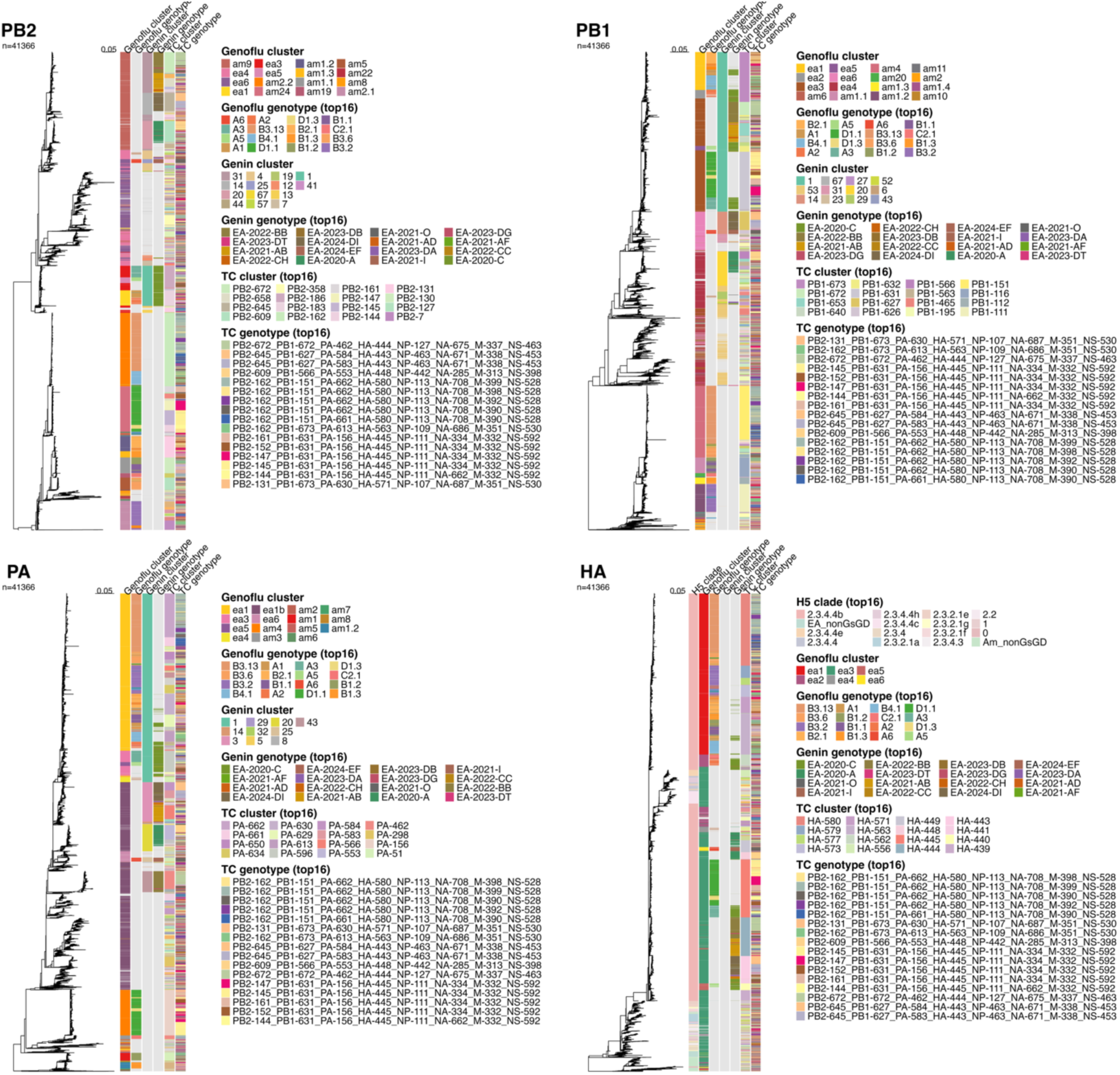
Maximum likelihood phylogenies for segments PB2, PB1, PA, and HA (n=41,366), comparing genotype and cluster labels for GenoFlu, Genin2, and TreeCluster (TC). HA also shows H5 clade (filled in using a phylogenetic neighbor algorithm). Top 16 most frequent labels were included in legends. Trees for NP, NA, M, and NS (Fig. S10).

We detected 39 Genin2 genotypes and three sub-genotypes within EA-2024-DI (DI.1, DI.2, and DI.2.1) in the dataset. Genin2 assigned fewer clusters per segment than GenoFlu (1–15 clusters per segment), with sizes from two tips up to 32,295, excluding M, which has a single cluster (size: 31,965) and HA, for which no clusters are assigned. Genin2 clusters were similarly non-monophyletic in several segments. For example, on the NS tree, Genin2 cluster 27 comprised 2472 sequences with purity of 6.1% (MRCA spanning 40,196 of 41,365 assigned tips), and cluster 29 comprised 589 sequences with purity of 1.4% (MRCA is the root), indicating how these sequences were interspersed throughout the tree.

TreeCluster inherently yields monophyletic clusters with 100% purity when phylogenies are treated as the ground truth. We evaluated multiple patristic distance thresholds and identified a pairwise patristic distance threshold of 0.04 subs/site as optimal, based on yielding a stable number of clusters (Fig. S6), a similar number of clusters in all segments (399–708 clusters per segment), minimal singletons (131–344 per segment), and a tolerable maximum cluster size of 6355 tips for PB2 (∼15% of the dataset; Fig. S6, Fig. S7). TreeCluster clusters showed clear separation of within- and between-cluster patristic distance distributions (Fig. S17). Segment clusters were concatenated into genotype constellations, resulting in 5042 unique genotypes, 2238 of which were observed in more than one sample (Fig. S21).

For CD-HIT, we evaluated cluster distributions for 85–99.5% genetic similarity (Fig. S8), across which we found 98% yielded a stable number of clusters at the elbow of the curve and a tolerable maximum cluster size (7692 for NS, ∼19% of the dataset). Despite partitioning the tree into a similar number of clusters (252–568 clusters per segment) and sizes as TreeCluster, CD-HIT clusters were not consistently monophyletic (Fig. 6). CD-HIT clusters had improved but incomplete separation of within- and between-clade patristic distance distributions (Fig. S16) compared to GenoFlu and Genin2. There were 6091 CD-HIT cluster constellations, of which 2186 were observed more than once.

### Diversity within predominant genotypes

We evaluated the TreeCluster genotypes and segment clusters comprising predominant GenoFlu and Genin2 genotypes (Fig. 5). Within GenoFlu genotype D1.1 (n=3525), we identified 21 TreeCluster genotypes (including singleton clusters), all due to variation in PB2 clusters (Table S5). All other segments had full agreement between GenoFlu and TreeCluster assignment of a single cluster within D1.1 (Fig. S11, Table S5). The span of D1.1 PB2 phylogenetic cluster IDs was close to the number of clusters; thus, contiguous cluster IDs are suggestive of PB2 diversification by drift. GenoFlu genotype D1.3 (n=352) shares six segments in common with D1.1 (PB1, PA, HA, NP, M, and NS), and its three PB2 clusters are a subset of D1.1’s PB2 clusters, indicating subdivision of a PB2 lineage rather than reassortment. The two genotypes differ at NA alone, consistent with D1.3 being a descendant reassortant of D1.1. The NA phylogenetic cluster carried by D1.3 (TreeCluster NA-662) was first detected in GenoFlu genotype A3 in North America sampled from bald eagles, glaucous gulls, and a red-tailed hawk in January 2022. The patristic distance comparison of D1.1 and D1.3 recovers this reassortment signal: the median between-genotype distance at NA was 0.345 subs/site versus 0.0071 subs/site within D1.1, a 48.4-fold separation, whereas every other segment separated by 1.5-fold or less, and the NA distance distributions did not overlap (minimum between-genotype distance 0.335 subs/site versus maximum within-D1.1 distance 0.028 subs/site). Additionally, if D1.1 and D1.3 sequences were hypothetically grouped together, the NA segment would have a span-to-cluster ratio of 164.5 while the PB2 segment would retain a ratio of 1.0 (despite comprising 20 distinct but contiguous clusters). The strong signal for NA reassortment based on the span-to-cluster ratio of D1.3 from D1.1 serves as a positive control for this approach.

**Figure 5.**
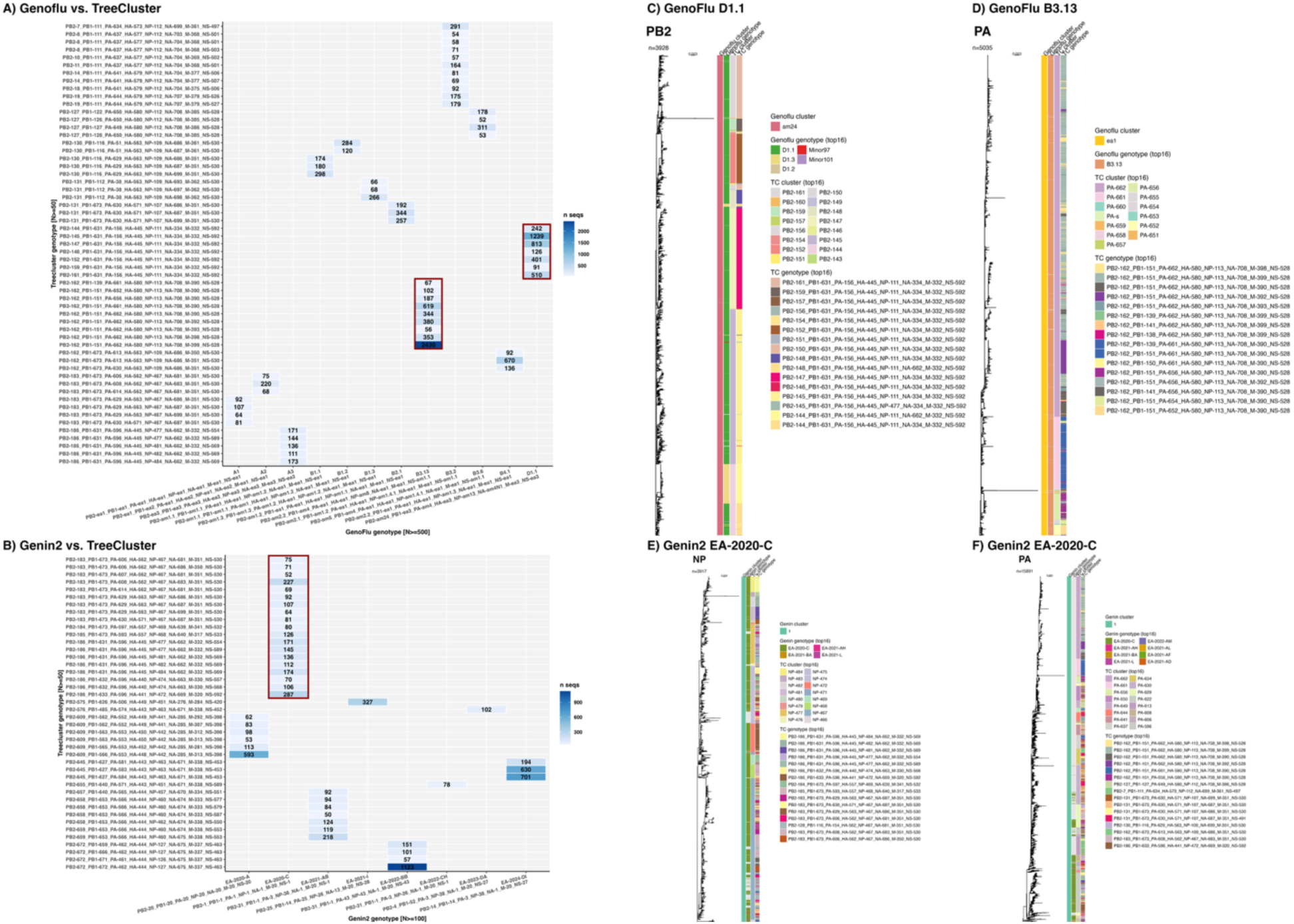
Phylogenetic diversity within predominant GenoFlu and Genin2 genotypes. Subtree cluster matrices of TreeCluster genotypes (with >50 samples) within predominant A) GenoFlu genotypes (with >500 samples), and B) Genin2 genotypes (with >100 samples). Red rectangles highlight cells for genotypes depicted in subtrees. Maximum likelihood phylogenies of segments with cluster diversity, clipped to subtrees at the most recent common ancestor of tips sharing genotype labels, for the C) PB2 tree for GenoFlu genotype D1.1, D) PA tree for GenoFlu genotype B3.13, and the E) NP and F) PA tree for Genin2 genotype EA-2020-C.

GenoFlu genotype B3.13 (n=4985) was broken into 85 TreeCluster genotypes (Table S5), 9 of which had at least 50 sequences (Fig. 5). B3.13 included diverse TreeCluster clusters: 22 PB1 clusters, 12 PA clusters, and 12 M clusters (Fig. S12), but their cluster ID spans were close to the number of clusters, suggestive of drift, not shift, in B3.13 (Table S5). The remaining segments for B3.13 agreed between GenoFlu and TreeCluster assignment of a single cluster. GenoFlu genotype B3.2 (n=2173) included 244 TreeCluster genotypes due to diverse clusters of PB2, PA, HA, NA, M, and NS. It displayed slightly wider cluster ID spans than expected due to drift (2.1 for M, 1.9 for NA, and 1.8 for HA).

B3.5 (n=153) showed wider than expected cluster ID spans in multiple segments: NS (ratio=10.7), M (6.7), PA (5.0), NA (4.5), and HA (3.5). Additionally, GenoFlu genotype A3 (n=971) had a wider NA span than expected (two clusters; ratio=9.0) and, to a lesser extent, NS (11 clusters; ratio=3.8). B4.1 (n=1121) and B3.1 (n=225) had moderately wider spans than expected for NA (both ratio=4.7), and B3.1 had a wider than expected HA cluster span, as well (ratio=3.2).

Some GenoFlu genotypes had full agreement with TreeCluster, including A6, C2.1, B3.3, and several minor genotypes (Table S5). Genotypes whose sequences fell within a single cluster for all segments acted as negative controls for this approach, demonstrating its ability to detect the absence of reassortment. GenoFlu genotype A6, for instance, comprises 315 sequences that share one TreeCluster genotype, so each segment contains a single cluster and the span and ratio are both 1.0, as expected for a genotype with no detected reassortment and minimal drift.

Genin2 genotypes also displayed diverse phylogenetic cluster combinations (Table S6). Within the largest Genin2 genotype, EA-2020-C (n=3361; Fig. 5; Fig. S13), no segment had full agreement for a single TreeCluster. Its segments comprised many clusters (seven PB2, 12 PB1, 30 PA, 19 NP, 27 NA, 32 M, and 29 NS). Notably, the span of TreeCluster IDs in EA-2020-C was wide for PB1 (12 clusters; span=521; ratio=43.4), and to some extent for NA (ratio=2.5) and NS (ratio=2.2), suggesting this genotype likely contains reassortants. The second largest Genin2 genotype, EA-2022-BB (n=1758; Fig. S14), comprised 60 TreeCluster genotypes made up of multiple clusters for PB2 (12 clusters), PB1 (18), PA (two), NP (11), and NS (three). However, none of their span-to-cluster ratios were suggestive of multiple reassortants. Other Genin2 genotypes with evidence of reassortment based on high span-to-cluster ratios include EA-2021-AB (n=1430; 17 PB1 clusters; span=173; ratio=10.2), and EA-2022-CH (n=150) for NS (three clusters; span=37; ratio=12.3) and PB1 (two clusters; span=14; ratio=7.0).

Among genotypes represented by at least two sequences, 39.4% of GenoFlu genotypes (26/66, representing 51.1% of GenoFlu-assigned sequences) and 33.3% of Genin2 genotypes (13/39, representing 62.9% of Genin2-assigned sequences), and were inconsistent with the phylogeny, having at least one segment with a span-to-cluster ratio of at least two. Genotypes made up of multiple adjacent TreeCluster clusters (ratio less than two in all segments) were considered consistent, as they reflect the finer resolution of phylogenetic clustering rather than incongruence.

### Comparison of cluster purity, monophyly, and phylogenetic diversity relative to TreeCluster

By design, TreeCluster yielded monophyletic clusters with 100% purity across all segments (Fig. 6) and the lowest median Faith’s PD in clusters (size>11, to account for differences in methods’ cluster size) in all segments (0.067–0.076 subs/site), and for size-matched clusters (Fig. S20). In comparison, GenoFlu clusters had lower median purity across segments, with PB1, HA, and NS as having the lowest purity (4.1%, 4.4%, 4.3%), and PB2 as having the highest (33.7%). As a result, a low proportion of GenoFlu clusters were monophyletic overall (14.2%) across segments (0% for HA, NA, M, and NS), with a maximum of 26.5% in PB2. The proportion of monophyletic GenoFlu clusters decreased with increasing cluster size (Fig. S18), going from 77.8% for clusters of size 2–5, down to 0% for clusters ≥ 10,000 tips. One large PB2 GenoFlu cluster was monophyletic (am5, 1039 tips), though eight larger PB2 clusters were not. Median Faith’s PD (cluster size>11) for GenoFlu ranged from 0.48 subs/site for PB1 to 6.05 subs/site for NA (Fig. S20).

**Figure 6.**
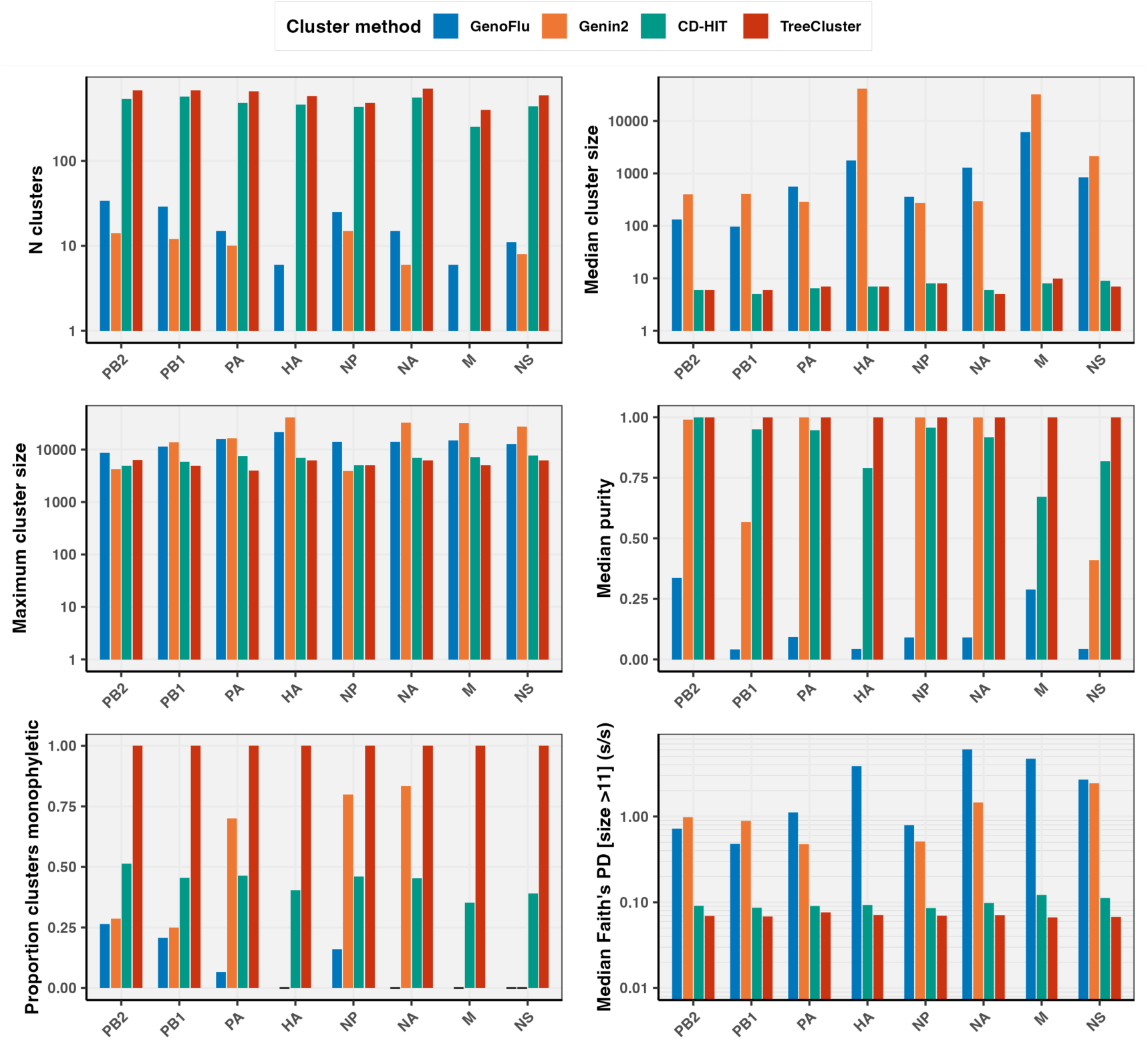
Comparison of H5Nx clustering methods’ cluster sizes, purity, monophyly, and phylogenetic diversity relative to TreeCluster. Panels from top-left, clockwise: number of clusters (log scale), median and maximum cluster size (log scale), median purity (tips assigned to cluster label of all tips in a clade descending from label’s MRCA), proportion clusters monophyletic (all descendants and the MRCA share cluster label and purity=1), and median Faith’s phylogenetic diversity (PD; minimum spanning tree) for cluster size greater than 11. Genin2 assigns a single M cluster, and no clusters for HA.

Genin2 clusters exhibited low monophyly for PB2, PB1, and NS (28.6%, 25.0%, and 0%) but higher monophyly for PA, NP, and NA (70.0%, 80.0%, and 83.3%), within the caveat that unassigned tips were ignored. Across all segments, 49.3% of all Genin2 clusters were monophyletic. Median purity in Genin2 clusters was lowest for NS and PB1 (40.9%, 56.6%); PA, NP, and NA were near 100% and PB2 was 99.0%. Genin2 does not assign HA clusters and assigns a single M cluster. Median Faith’s PD (for cluster size>11) for Genin2 clusters overall ranged from 0.47 (PA) to 2.44 (NS) subs/site.

At the chosen thresholds, CD-HIT yielded a comparable number of clusters as TreeCluster (across segments: 252–568 versus 399–708) and median cluster size (5–9 versus 5–10), but lower monophyly (35.3–51.3% versus 100%). Overall, 44.4% of clusters were monophyletic. However, CD-HIT purity was moderate to high (median 67.1–100% across segments, overall median 90.2%), which was closer to Genin2 and TreeCluster than to GenoFlu (median purity 4–34% across segments). CD-HIT purity was highest in PB2 (100%), NP (95.8%), PB1 (94.9%), PA (94.6%), and NA (91.7%), but lower in M (67.1%), HA (79.1%), and NS (81.8%), possibly indicative of higher homoplasy. The median Faith’s PD (for cluster size>11) for CD-HIT across segments (0.086–0.122 subs/site) was closer to TreeCluster than to GenoFlu or Genin2. At matched cluster size, GenoFlu, Genin2, and CD-HIT clusters generally had higher median PD. For cluster sizes > 200, TreeCluster’s median Faith PD (0.32 subs/site) was two-to eight-fold lower than the other methods (Fig. S20).

## Discussion

Downstream research and public health applications of H5Nx genomes often rely on the assumption that genotype labels, such as those assigned by GenoFlu and Genin2, delineate sets of genomes with recent shared ancestry. Using 41,366 globally sampled H5Nx genomes, we characterised the limitations of genetic similarity-based approaches and illustrated the contexts in which GenoFlu and Genin2 genotypes may not be reliable. Despite providing rapid, relatively stable, and easily communicated labels for broad outbreak surveillance, these genetic similarity-based genotyping approaches failed to assign sequences divergent from their reference sets, grouped genomes that do not share recent common ancestry, and obscured mutational drift and reassortment within major circulating genotypes. While we did not demonstrate the superiority of phylogenetic-clustering using TreeCluster *per se* (given the circularity of evaluating phylogenetic clusters against the same phylogenies from which they are derived) these findings suggest that a phylogenetically-based genotyping system may facilitate more effective evolutionary analyses in response to the H5 clade 2.3.4.4b panzootic and to IAV broadly.

Neither GenoFlu nor Genin2 could describe the global diversity of H5Nx, with only 63.8% of 2020-2025 sequences assigned genotypes by GenoFlu and 34.4% by Genin2. This was expected as these tools were only designed to assign 2.3.4.4b genotypes within their respective focal regions. However, even within these regions, genotype assignment remained incomplete (2.6% of 2020-2025 North American sequences unassigned by GenoFlu; 3.1% of European sequences assigned by Genin2) despite our relatively stringent genome quality control and the sample collection date cut-off preceding the release date of the evaluated tool versions. This incomplete assignment, with notable segment-level variability (Fig. 3), reflects the limitations of genetic similarity-based systems being reliant on a fixed set of reference sequences that may not fully capture circulating diversity. However, non-assignment was not only linked with the emergence of novel diversity: TreeCluster genotype alias TC4771 comprised 2430 sequences assigned to GenoFlu genotype B3.13 together with 20 sequences GenoFlu left unassigned, which were phylogenetically indistinguishable from the rest of the cluster. Interestingly, high assignment rates outside a tool’s focal region (such as GenoFlu’s strong performance in South America and Antarctica, or Genin2’s assignments in Western and Middle Africa) highlighted known epidemiological and flyway connections, as well as differences in segments’ conservation. The practical consequence of this reference dependence in genetic similarity-based typing is that genotype-based surveillance summaries, or any analysis that drops unassigned sequences, risks being biased toward well-represented regions, hosts, and viral diversity. This presents a major challenge when attempting to investigate under-sampled geographies, new spillovers, and rapidly diversifying lineages.

Even when sequences were similar enough to references to be assigned genotypes, the underlying segment clusters frequently grouped genetically similar segments that were not related by recent shared ancestry. GenoFlu clusters exhibited low median purity (ranging from 4.1% for PB1 to 33.7% for PB2), with only 14.2% of clusters being monophyletic across all segments. Genin2 clusters were more commonly monophyletic (49.3%) and achieved higher median purity (nearly 100% for PA, NP, PB2, and NA, but 40.9% for NS). While this may reflect the initial phylogenetic basis of Genin2 training clusters (among other implementation differences), this comparison is also biased in Genin2’s favor as unassigned tips were excluded when calculating purity and Genin2 assigned roughly half as many sequences as GenoFlu. It should also be noted that not all non-monophyly carries the same consequences. Paraphyly (the MRCA is likely a member of the cluster, but not all descendants are members) due to the presence of nested clusters is expected as a clade diversifies into subclades, however, it reflects a violation of the recent shared ancestry assumption, in addition to instances where clusters are polyphyletic (the MRCA is not likely in the cluster, nor are all the descendants) due to a cluster label appearing to emerge multiple times independently across clades. Monophyly and purity declined as cluster size increased for GenoFlu and Genin2, however, having larger clusters did not fully account for these tools’ suboptimal purity. When we applied CD-HIT (at a 98% genetic similarity threshold), it generated a similar number and size of clusters as TreeCluster (at a 0.04 subs/site tree distance threshold), yet only 44% of CD-HIT clusters were monophyletic with low median purity in M (67.1%), HA (79.1%), and NS (81.8%). This may reflect higher rate variation or homoplasy in shorter and surface protein segments, corresponding to less well-resolved phylogenies. However, given the consistently lower cluster purity, reduced monophyly, and poor separation of within-versus between-cluster patristic distances observed across all similarity-based methods, this should be mainly attributed to the limitations of assigning clusters by genetic similarity.

We next asked whether the distribution of phylogenetic clusters within a genotype could distinguish within-lineage diversification (drift) from reassortment (shift). A genotype that has accumulated mutations within a lineage would be expected to occupy contiguous TreeCluster cluster IDs and therefore the numerical span of cluster IDs relative to the number of clusters would be near 1.0. On the other hand, a reassortant genotype that combines clusters from distant parts of their respective segment trees should yield an inflated span-to-cluster ratio. Genotypes confined to one TreeCluster cluster per segment (e.g., GenoFlu A6 [n=315]) trivially returned a span-to-cluster of 1.0. Conversely, if we pooled GenoFlu genotypes D1.1 and D1.3, which differ primarily through reassortment of NA, the NA segment yielded a high span-to-cluster ratio (164.5) while PB2 and other segments’ ratios remained at 1. This suggests that this heuristic may be able to isolate reassortment to discordant segments without being confounded by within-segment diversification. Furthermore, this analysis also traced the donor NA segment of D1.3 (TreeCluster NA-662) to GenoFlu A3 viruses circulating in wild birds since early 2022. However, as cluster IDs are assigned in ladderized tip order this heuristic is sensitive to the tip order and would not be invariant under alternative ladderizations. Incongruence testing with TreeSort (Markin et al., 2025) would be the natural approach to corroborate this metric, but it presents computational challenges with phylogenies of large scale. Similarly, integration of bootstrap support, formal permutation testing, and validation across broader IAV diversity are needed to formalise this heuristic.

Nevertheless, applying this heuristic to predominant H5Nx genotypes revealed otherwise hidden diversity due to drift and reassortment. For example, we observed pronounced mutational drift (indicated by multiple contiguous TreeCluster clusters with span ratios near 1.0) in PB2 within D1.1 and D1.3; in PB1, PA, and M within the dairy cattle-associated B3.13; and across six segments in B3.2. For D1.1, our findings corroborate the rapid diversification of PB2 via drift following its initial emergence by reassortment (Crespo-Bellido et al., 2025). This sort of sublineage diversification may warrant finer reference partitioning and targeted evaluation for phenotypic or antigenic divergence. More concerningly, our analysis also found evidence of potential cryptic reassortment (substantially inflated cluster ID spans) within GenoFlu genotypes B3.5, A3, B4.1, and B3.1, and for Genin2 genotypes EA-2020-C, EA-2021-AB, and EA-2022-CH. When a single genotype label inadvertently combines distinct reassortants, downstream analyses that treat the genotype as a single evolutionary unit or concatenate segment alignments can be misleading due to the incongruence of the underlying phylogenies (Dugan et al., 2008). In phylodynamics and phylogeographic modeling, this can bias estimates of viral emergence dates, evolutionary rate, host transition, and spatial diffusion.

H5Nx clustering and genotyping approaches present trade-offs that are often driven by the motivation for their development. For example, GenoFlu and Genin2 provide a relatively concise number of labels that are effective for high-level communication around outbreak response. Their use of fixed references also enables computationally efficient assignments that are unlikely to drastically change over time as tools are updated. Phylogenetic clustering, by contrast, explicitly accommodates deviations of genetic and phylogenetic distance due to homoplasy, reversions, and site and branch-level evolutionary rate variation, but introduces uncertainty associated with sequence alignment, substitution model assumptions, and tree topology. These sources of uncertainty can be amplified by low-quality sequences, temporal outliers, long branches, and dense sampling of highly similar sequences. Although these can be mitigated by quality control measures, as performed here, this uncertainty also means phylogenetic cluster assignments are inherently less stable than reference-based classification. Incorporating new sequences can change the topology and branch lengths of the underlying phylogenies which can shift cluster boundaries and lead to clusters merging or splitting. Future work should directly evaluate the stability of H5Nx phylogenetic genotype designation in response to random and biased subsampling, holding out dates, or new data, as well as evaluate the possibilities of bootstrap support values to handle phylogenetic uncertainty.

Other phylogenetically-informed clustering and lineage designation approaches have been proposed for IAV, but they have some limitations for H5Nx. An influenza nomenclature framework was developed to identify HA and NA subclades based on the group size, distance from the parent, and amino acid substitutions on the branch preceding the group’s common ancestor (Neher et al., 2026). This approach is well-suited to human seasonal influenza viruses, where clade names are antigenically and epidemiologically meaningful, and though it could be expanded for all eight segments of H5Nx, it may need consideration of sampling biases related to hosts and geographies. Similarly, multidimensional scaling with k-means clustering using evolutionary distance has also been applied to identify H5N1 clusters (Tawidian et al., 2024), however it is unclear how this approach would compare to phylogenetic clustering, especially in terms of handling convergent evolution.

By adapting recently developed genotyping approaches for non-segmented viruses, it may be possible to develop a practical H5Nx genotyping method that retains evolutionary fidelity while achieving the computational efficiency, stability, and label clarity of genetic similarity-based approaches. For example, to avoid expensive *de novo* reconstruction of phylogenies, a set of carefully updated global phylogenies could be maintained for each segment and new genome sequences placed into these trees using maximum parsimony using UShER (Turakhia et al., 2021) or TIPars (Ye et al., 2024). To achieve label stability, an operational H5Nx nomenclature could anchor phylogenetic cluster names to a central or early representative sequence, analogous to approaches used by HIV phylogenetic cluster monitoring programs (Poon et al., 2015; Poon et al., 2016; McLaughlin et al., 2021; Miller et al., 2022; McLaughlin et al., 2025). The Pango system of hierarchical lineage nomenclature, used in SARS-CoV-2 (Rambaut et al., 2020; O’Toole et al., 2021), Mpox virus (Happi et al., 2022; Ruis et al., 2025), rabies virus (Campbell et al., 2022), human respiratory syncytial virus (Goya et al., 2024), dengue virus (Hill et al., 2024), and chikungunya virus (Colquhoun et al., 2026), could also be adopted to provide a compact labeling system for tracking related groups while accommodating emerging diversity. Segment lineages could be assigned separately and then combined into epidemiologically meaningful lineage constellations and assigned alphanumeric genotype aliases. Finally, tools such as Autolin (McBroome et al., 2024), which use a genotype representation index to identify nodes that represent distinct, compact, and sufficiently large groups, could be applied to each segment tree to support automated genotype designations.

This study has several limitations. We relied upon publicly available data that are affected by sampling biases related to outbreak severity, host mortality and survival, access to samples, geography, sequencing capacity, and the proximity or importance of hosts to humans. Further sampling bias may have been imparted by filtering to whole genomes, which excluded partial sequences from tree inference. Together these sampling biases impact observed distributions of diversity, cluster size, and geographies. Additional challenges and limitations common to all genotyping approaches include the trade-off of sensitivity and specificity, effective parameterisation, and appropriate usage. Genotyping trades flexibility to accommodate novel diversity against sensitivity and stability of established lineages. For example, LABEL, a hidden Markov model using support vector machines for H5N1 lineage assignment (Shepard et al., 2014), struggled to identify lineages divergent from references (Turakhia et al., 2021). Approaches are sensitive to their parameterisations (including but not limited to distance thresholds), yet optimal parameter values depend on viruses’ natural history (such as sampling rate, transmission routes, bottlenecks, duration of infectiousness, and strength of within-host selection) and intended downstream applications. For example, despite similar evolutionary clock rates for HIV and IAV (∼10⁻³ substitutions/site/year) (Chen and Holmes 2006; Vrancken et al., 2014), HIV-1 subtype B phylogenetic clustering in British Columbia utilizes a threshold of 0.02 subs/site to identify epidemiologically-linked cases (Poon et al., 2015; Poon et al., 2016; McLaughlin et al., 2021; Miller et al., 2022; McLaughlin et al., 2025); whereas, we selected a threshold of 0.04 subs/site for H5Nx to yield evenly sized clusters. No genotyping system is optimised for all possible downstream applications. An additional limit to this analysis is that we did not evaluate how the proposed framework would perform across broader IAV diversity. We have applied TreeCluster to H3 clustering (*Tran et al., 2026, in preparation*), however, a global IAV analysis would better illuminate the frequency and origins of reassortment. Future work to characterize IAV diversity could also consider minority allele populations from raw reads, as the presence of mixed infections and minority variants are not revealed through consensus sequences, unless multiple HA and NA types are reported. It is possible that some of the cluster constellations we identified are artefacts of mixed infections or defective reassortants, however, the detection of a constellation in multiple samples supports its viability for onward transmission.

## Conclusion

Viral classification is critical for genomic surveillance of H5Nx and other IAV, yet designation and assignment of genotypes based on genetic similarity to reference sequences does not reliably yield clusters that are congruent with global phylogenies. While genetic similarity clustering has the advantages of speed, scalability, and stability, and suffices for some downstream applications, phylogenetic clustering offers granularity to resolve combinations of H5Nx clusters across the genome that were not captured by sequence similarity-based approaches, highlighting genotypes with evidence of grouping multiple reassortants. Phylogenetic inference provides evolutionary context to define groups of viruses that share recent ancestry, as is necessary for some downstream approaches including the identification of novel reassortants and phylodynamic inference. Furthermore, genotype surveillance and modeling rely upon representative data and references across geographies, hosts, and viral diversity. Further incorporating phylogenetic relationships into H5Nx genotyping could improve surveillance, situational awareness, modeling, and risk assessment as H5Nx and IAV continues to diversify.

## Supporting information

GISAID supplementary file

NCBI Virus supplementary file

## Acknowledgments

We gratefully acknowledge all originating and submitting laboratories of H5Nx genome sequences and metadata on GISAID and NCBI. We are grateful for feedback received at the One Health Antimicrobial Resistance and Emerging Zoonoses meeting, Dynamics and Evolution of Human Viruses conference, and the Symposium for the Canadian Society for Virology. Generative AI (Claude Code, Anthropic, Opus 5.5) was used for de-bugging, testing code, modifying figures, and expanding comments. The authors reviewed all code and outputs, and take full responsibility for code, analyses, results, and interpretations.

## Data availability

Access to sequences originating from GISAID is limited to approved data users. All GISAID genome sequences and associated metadata supporting the findings of this study can be accessed at the EpiSet digital object identifier, doi.org/10.55876/gis8.260622ts. The NCBI Virus accession IDs used in the final cleaned dataset are provided in supplementary file, NCBI_accessions.csv. Code, summary tables, and plots are available on GitHub (https://github.com/angmcl/h5nx_phylo_geno) and archived on Zenodo (https://doi.org/10.5281/zenodo.23606951).

