## Supplementary Materials for "Influenza A virus H5Nx genotypes assigned by genetic similarity are incongruent with global phylogenies"

**Supplementary Tables and Figures**

**Table S1.** Gap-trimming nucleotide alignments of eight H5Nx segments.

**Table S2.** Exclusion of temporal outliers from preliminary phylogenies.

**Table S3.** List of all unique common hosts and lowest taxonomic level in the cleaned H5Nx dataset.

**Table S4.** Countries with data in global regions within the cleaned and de-duplicated dataset.

**Table S5.** Phylogenetic cluster composition of GenoFlu genotypes with at least 10 sequences.

**Table S6.** Phylogenetic cluster composition of Genin2 genotypes with at least 10 sequences.

**Fig. S1.** Identification and removal of temporal outliers, shown for HA segment.

**Fig. S2.** Cleaned and de-duplicated H5Nx samples for the full date range 1961-2025 (n=41,366).

**Fig. S3.** The distribution of standardized common hosts in the dataset.

**Fig. S4.** Maximum likelihood HA phylogeny colored by H5 clade before and after filling in labels using a phylogenetic neighbourhood algorithm.

**Fig. S5.** Maximum likelihood NA phylogeny colored by NA type before and after filling in labels using a phylogenetic neighbourhood algorithm.

**Fig. S6.** Evaluation of TreeCluster maximum pairwise patristic distance across segments.

**Fig. S7.** Phylogenetic cluster size distribution with TreeCluster threshold: 0.04 subs/site.

**Fig. S8.** Evaluation of CD-HIT clusters at 85-99% minimum genetic similarity across segments.

**Fig. S9.** Genetic cluster size distribution from CD-HIT with 98% similarity threshold.

**Fig. S10**. Maximum likelihood phylogenies for segments NA, NP, M, and NS (n=41,366) comparing genotype and cluster labels for Genoflu, Genin2, and TreeCluster (TC).

**Fig. S11.** Maximum likelihood trees for GenoFlu genotype D1.1.

**Fig. S12.** Maximum likelihood trees for GenoFlu genotype B3.13.

**Fig. S13.** Maximum likelihood trees for Genin2 genotype EA-2020-C

**Fig. S14.** Maximum likelihood trees for Genin2 genotype EA-2022-BB

**Fig. S15.** Distributions of pairwise patristic distances (subs/site) within or between GenoFlu clusters

**Fig. S16.** Distributions of pairwise patristic distances within or between CD-HIT clusters (98%).

**Fig. S17.** Distributions of pairwise patristic distances within or between TreeClusters (0.04 subs/site).

**Fig. S18.** Proportion of monophyletic clusters across cluster size, method, and segment.

**Fig. S19.** Clusters’ median purity across cluster size, method, and segment.

**Fig. S20**. Clusters’ median Faith’s phylogenetic diversity (PD) cluster size, method, and segment.

**Fig. S21.** H5Nx TreeCluster genotypes connect unique combinations of segment clusters.

**Fig. S22**. Phylogenetic cluster span ratio within predominant genotypes to distinguish drift from reassortment

**Table S1.** Gap-trimming nucleotide alignments for eight H5Nx segments. Initial and trimmed widths of nucleotide alignments, and number and proportion of positions trimmed. Ends were trimmed to the first and last positions with ≤90% gaps (minimum nucleotide end: 0.1). Internal positions were trimmed if they exceeded 99% gaps (retaining insertions/deletions as infrequent as 1% of sequences). Number of sequences precedes quality filtering, de-duplication by host, and removal of temporal outliers.

| **Segment** | **N seqs** | **Width initial (b)** | **Width trimmed (b)** | **Width removed (b)** | **Proportion of positions removed** | **Minimum nucleotide end** |
| --- | --- | --- | --- | --- | --- | --- |
| PB2 | 61,491 | 4700 | 2341 | 2359 | 0.50 | 0.1 |
| PB1 | 61,491 | 4787 | 2342 | 2445 | 0.51 | 0.1 |
| PA | 61,491 | 2975 | 2233 | 742 | 0.25 | 0.1 |
| HA | 61,491 | 2222 | 1776 | 446 | 0.20 | 0.1 |
| NP | 61,491 | 2896 | 1566 | 1330 | 0.46 | 0.1 |
| NA | 61,491 | 3836 | 1588 | 2248 | 0.59 | 0.1 |
| M | 61,491 | 1784 | 1027 | 757 | 0.42 | 0.1 |
| NS | 61,491 | 2527 | 890 | 1637 | 0.65 | 0.1 |

**Table S2.** Exclusion of temporal outliers from preliminary phylogenies. For each segment FastTree (N tip initial), residuals were calculated from the root-to-tip (RTT) linear regression. Residual outliers were removed if residuals exceeded the maximum residual (10 standard deviations). Pendant edge outliers were removed if mutations on the terminal branch exceeded the maximum mutations (6% of alignment width). Samples were excluded if any segment sequences were excluded to yield N tip final.

| **Segment** | **N tip**  **initial** | **Trimmed alignment (b)** | **Maximum residual** | **N residuals outliers** | **Maximum mutations** | **Pendant edge outliers** | **N tip**  **final** |
| --- | --- | --- | --- | --- | --- | --- | --- |
| PB2 | 41,427 | 2341 | 0.511 | 3 | 140 | 17 | 41,366 |
| PB1 | 41,427 | 2342 | 0.237 | 4 | 141 | 8 | 41,366 |
| PA | 41,427 | 2233 | 0.210 | 4 | 134 | 10 | 41,366 |
| HA | 41,427 | 1776 | 0.447 | 3 | 107 | 7 | 41,366 |
| NP | 41,427 | 1566 | 0.261 | 4 | 94 | 3 | 41,366 |
| NA | 41,427 | 1588 | 0.998 | 3 | 95 | 18 | 41,366 |
| M | 41,427 | 1027 | 0.231 | 4 | 62 | 4 | 41,366 |
| NS | 41,427 | 890 | 0.523 | 4 | 53 | 4 | 41,366 |

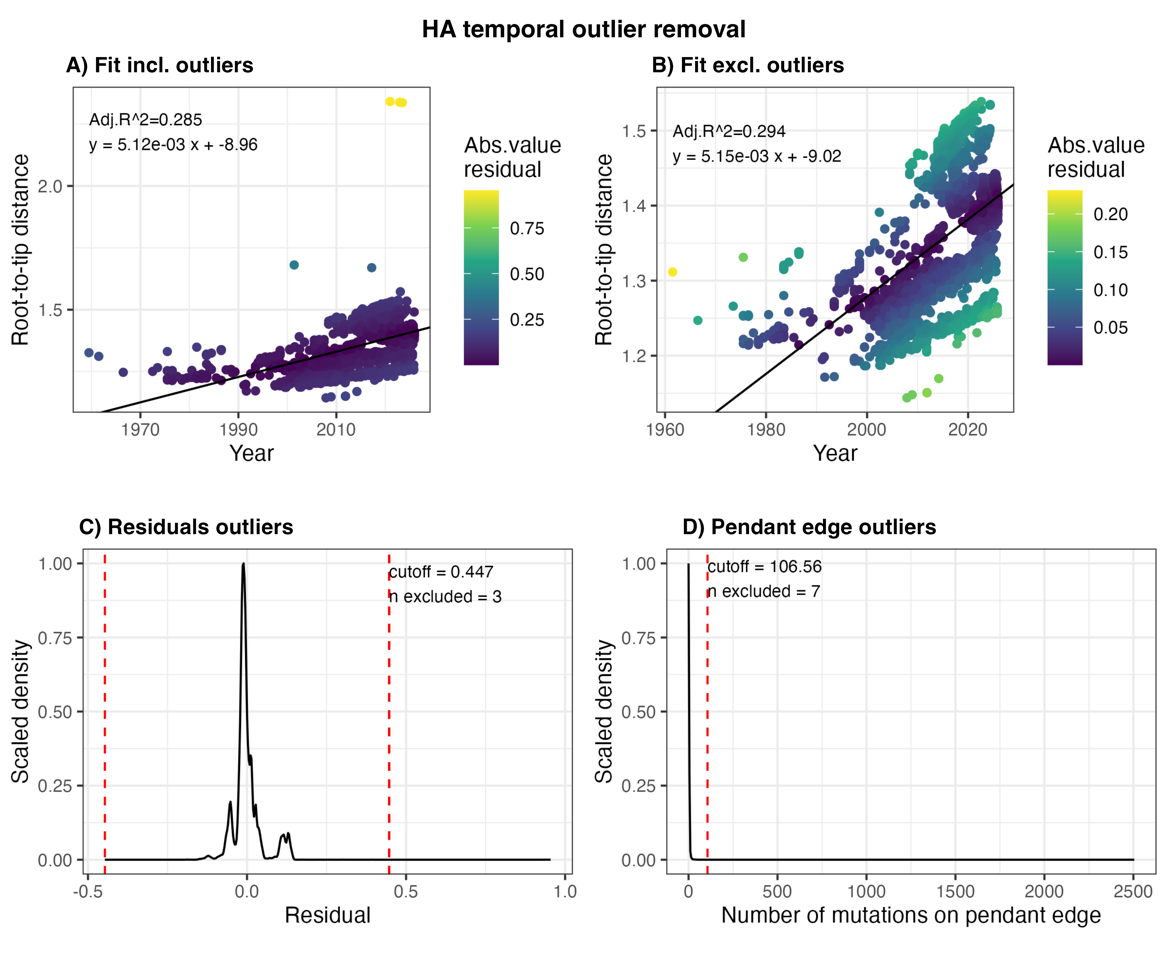
**Fig. S1.** Identification and removal of temporal outliers on preliminary trees, shown for HA segment. A) Root-to-tip (RTT) regression including and B) excluding outliers, annotated with adjusted R-squared and fit, coloured by absolute value residual. C) Residuals outliers were > 10 standard deviations from the mean residual. D) Pendant edge outliers had substitutions exceeding 6% of the trimmed alignment width.

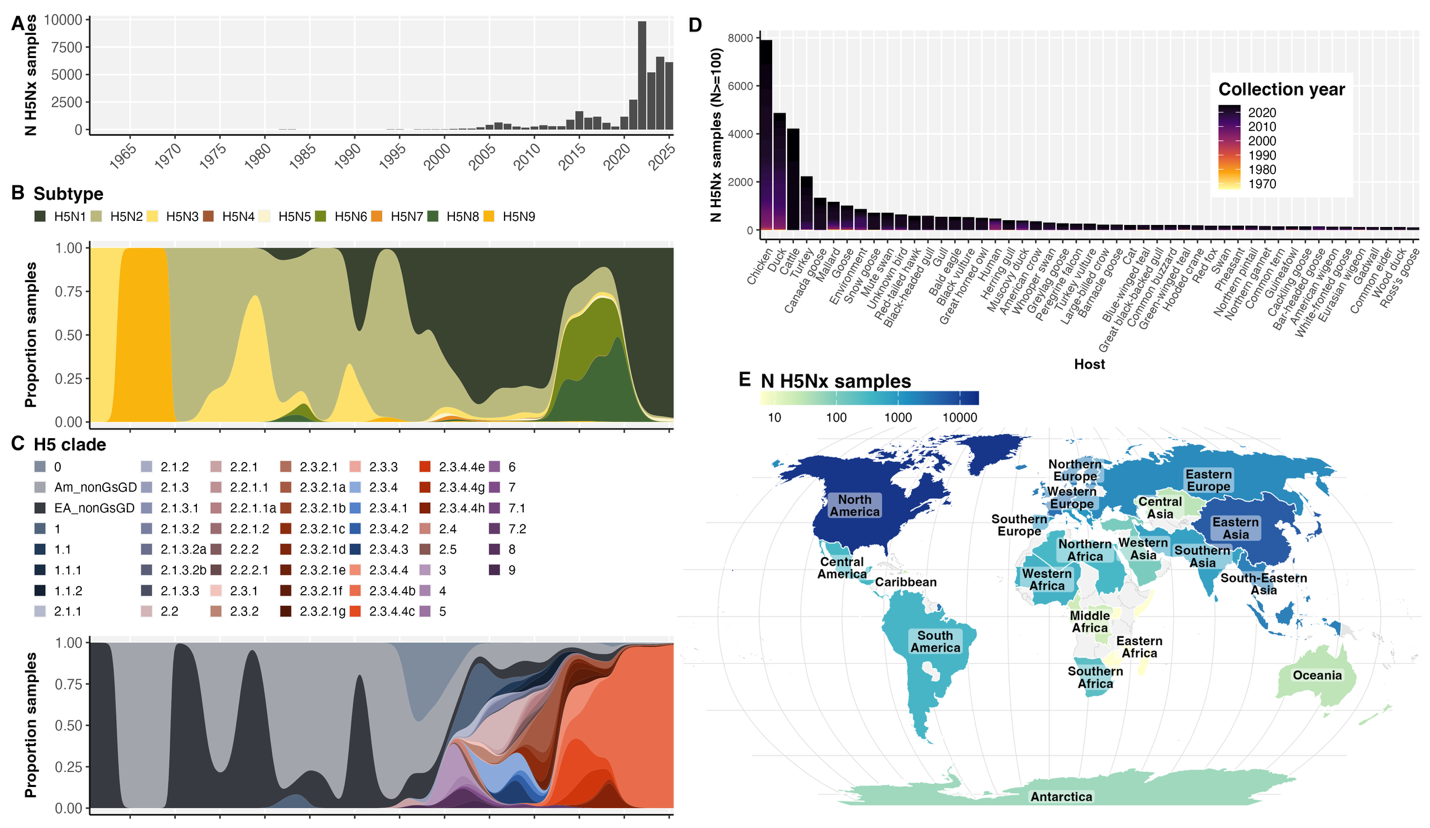
**Fig. S2.** Cleaned and de-duplicated H5Nx samples for the full date range 1961-2025 (n=41,366). A) Distribution of clean H5Nx samples across collection years; proportion of samples by B) subtypes and C) H5 clade; D) hosts with at least 100 samples (complete set in Fig. S3); and E) number of samples available from global regions.

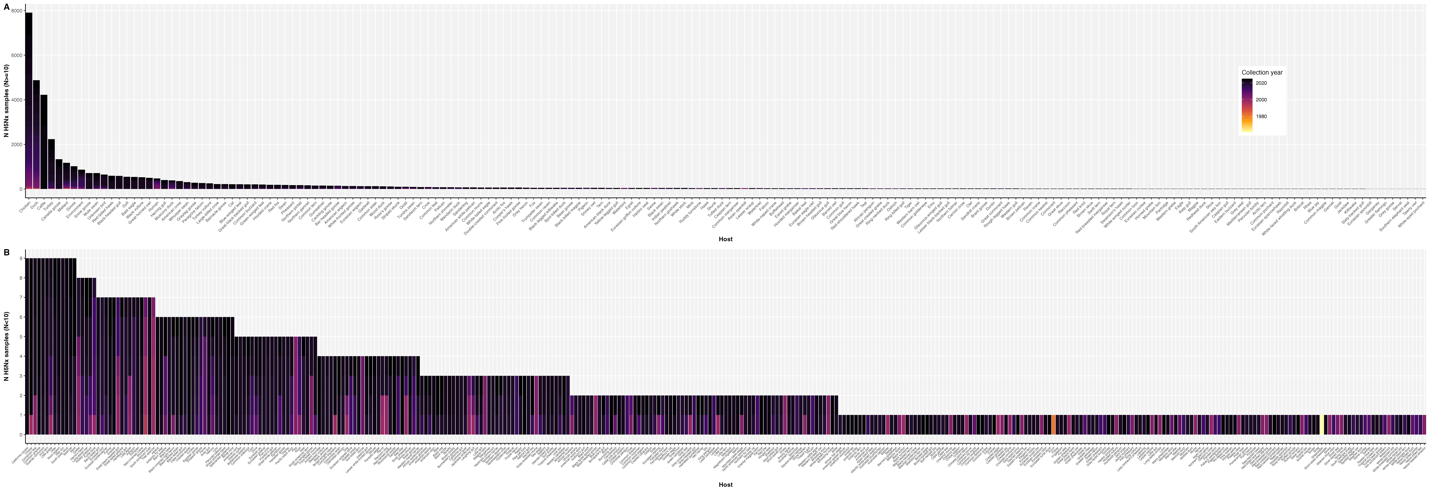
**Fig. S3.** The distribution of standardized common hosts in the dataset. Common names of hosts with A) at least 10 samples, B) and fewer than 10 samples over the full date range. Common names were spell-checked and standardized to the most specific common name.

**Table S3.** List of all unique common hosts and lowest taxonomic level in the cleaned H5Nx dataset. Sorted alphabetically by common host (*Genus species* if available, else lowest taxonomic level). Excludes samples from environment, food, and dairy products.

African fish eagle (*Haliaeetus vocifer*), African houbara (*Chlamydotis undulata*), African jacana (*Actophilornis africanus*), African oystercatcher (*Haematopus moquini*), African penguin (*Spheniscus demersus*), Alpaca (*Vicugna pacos*), American black duck (*Anas rubripes*), American coot (*Fulica americana*), American crow (*Corvus brachyrhynchos*), American golden-plover (*Pluvialis dominica*), American goshawk (*Astur atricapillus*), American kestrel (*Falco sparverius*), American robin (*Turdus migratorius*), American white pelican (*Pelecanus erythrorhynchos*), American wigeon (*Mareca americana*), American wood stork (*Mycteria americana*), American woodcock (*Scolopax minor*), Anatinae (Anatidae), Ancient murrelet (*Synthliboramphus antiquus*), Antarctic fur seal (*Arctocephalus gazella*), Antarctic shag (*Leucocarbo bransfieldensis*), Antarctic tern (*Sterna vittata*), Arctic fox (*Vulpes lagopus*), Arctic skua (*Stercorarius*), Arctic tern (*Sterna paradisaea*), Armenian gull (*Larus armenicus*), Asian black bear (*Ursus thibetanus*), Asian openbill (*Anastomus oscitans*), Atlantic puffin (*Fratercula arctica*), Atlantic white-sided dolphin (*Leucopleurus acutus*), Audouin's gull (*Ichthyaetus audouinii*), Australian brushturkey (*Alectura lathami*), Australian pelican (*Pelecanus conspicillatus*), Babbler (Timaliidae), Badger (*Meles*), Baikal teal (*Sibirionetta formosa*), Bald eagle (*Haliaeetus leucocephalus*), Band-tailed gull (*Larus belcheri*), Bar-headed goose (*Anser indicus*), Barn swallow (*Hirundo rustica*), Barnacle goose (*Branta leucopsis*), Barred owl (*Strix varia*), Barrow's goldeneye (*Bucephala islandica*), Bean goose (*Anser fabalis*), Bear (*Ursus*), Bearded vulture (*Gypaetus barbatus*), Bewicks swan (*Cygnus bewickii*), Black bear (*Ursus americanus*), Black bulbul (*Hypsipetes leucocephalus*), Black kite (*Milvus migrans*), Black scoter (*Melanitta americana*), Black skimmer (*Rynchops niger*), Black swan (*Cygnus atratus*), Black swift (*Cypseloides niger*), Black turnstone (*Arenaria melanocephala*), Black vulture (*Coragyps atratus*), Black-billed magpie (*Pica hudsonia*), Black-crowned night heron (*Nycticorax nycticorax*), Black-faced spoonbill (*Platalea minor*), Black-headed goose (Anatidae), Black-headed gull (*Chroicocephalus ridibundus*), Black-headed heron (*Ardea melanocephala*), Black-legged kittiwake (*Rissa tridactyla*), Black-necked crane (*Grus nigricollis*), Black-necked swan (*Cygnus melancoryphus*), Black-tailed gull (*Larus crassirostris*), Blackish oystercatcher (*Haematopus ater*), Blakiston's fish owl (*Bubo blakistoni*), Blowfly (*Calliphora*), Blue crane (*Anthropoides paradiseus*), Blue jay (*Cyanocitta cristata*), Blue-winged teal (*Spatula discors*), Bobcat (*Lynx rufus*), Bonaparte's gull (*Chroicocephalus philadelphia*), Booted eagle (*Hieraaetus pennatus*), Bottlenose dolphin (*Tursiops*), Brant goose (*Branta bernicla*), Broad-winged hawk (*Buteo platypterus*), Brown booby (*Sula leucogaster*), Brown pelican (*Pelecanus occidentalis*), Brown skua (*Stercorarius lonnbergi*), Brown-headed gull (*Chroicocephalus brunnicephalus*), Brown-hooded gull (*Chroicocephalus maculipennis*), Buff-necked ibis (*Theristicus caudatus*), Bufflehead (*Bucephala albeola*), Bulbul (Pycnonotidae), Burmeister's porpoise (*Phocoena spinipinnis*), Burrowing owl (*Athene cunicularia*), Bush dog (*Speothos venaticus*), Cackling goose (*Branta hutchinsii*), California condor (*Gymnogyps californianus*), California gull (*Larus californicus*), Canada goose (*Branta canadensis*), Canvasback (*Aythya valisineria*), Cape cormorant (*Phalacrocorax capensis*), Cape gannet (*Morus capensis*), Cape teal (*Anas capensis*), Carrion crow (*Corvus corone*), Caspian gull (*Larus cachinnans*), Caspian tern (*Hydroprogne caspia*), Cat (*Felis catus*), Cattle (*Bos taurus*), Chestnut munia (*Lonchura atricapilla*), Chestnut teal (*Anas castanea*), Chicken (*Gallus gallus*), Chilean dolphin (*Cephalorhynchus eutropia*), Chiloe wigeon (*Mareca sibilatrix*), Chimango (*Milvago chimango*), Chinese pond heron (*Ardeola bacchus*), Chinstrap penguin (*Pygoscelis antarcticus*), Chukar (*Alectoris chukar*), Cinnamon teal (*Spatula cyanoptera*), Civet (*Civettictis civetta*), Collared plover (*Anarhynchus collaris*), Common blackbird (*Turdus merula*), Common buzzard (*Buteo buteo*), Common crane (*Grus grus*), Common dolphin (*Delphinus delphis*), Common eider (*Somateria mollissima*), Common goldeneye (*Bucephala clangula*), Common grackle (*Quiscalus quiscula*), Common gull (*Larus canus*), Common hill myna (*Gracula religiosa*), Common kestrel (*Falco tinnunculus*), Common kingfisher (*Alcedo atthis*), Common loon (*Gavia immer*), Common magpie (*Pica pica*), Common merganser (*Mergus merganser*), Common moorhen (*Gallinula chloropus*), Common murre (*Uria aalge*), Common pheasant (*Phasianus colchicus*), Common pochard (*Aythya ferina*), Common quail (*Coturnix coturnix*), Common raven (*Corvus corax*), Common redshank (*Tringa totanus*), Common shelduck (*Tadorna tadorna*), Common snipe (*Gallinago gallinago*), Common teal (*Anas crecca*), Common tern (*Sterna hirundo*), Common wood pigeon (*Columba palumbus*), Cooper's hawk (*Accipiter cooperii*), Coot (*Fulica*), Cormorant (Phalacrocoracidae), Coscoroba swan (*Coscoroba coscoroba*), Coyote (*Canis latrans*), Cranes (*Grus*), Crested auklet (*Aethia cristatella*), Crested caracara (*Caracara plancus*), Crested eagle (*Morphnus guianensis*), Crested goshawk (*Accipiter trivirgatus*), Crested jay (*Platylophus galericulatus*), Crested myna (*Acridotheres cristatellus*), Crow (*Corvus*), Crowned cormorant (*Microcarbo coronatus*), Curlew (*Numenius*), Dalmatian pelican (*Pelecanus crispus*), Dark-eyed junco (*Junco hyemalis*), Daurian starling (*Agropsar sturninus*), Dog (*Canis lupus*), Dolphin (*Tursiops*), Double-crested cormorant (*Phalacrocorax auritus*), Dove (Columbidae), Duck (Anatidae), Dunlin (*Calidris alpina*), Eagle (*Aquila*), Eared grebe (*Podiceps nigricollis*), Eastern buzzard (*Buteo japonicus*), Eastern screech owl (*Megascops asio*), Egret (*Ardea alba*), Egyptian goose (*Alopochen aegyptiaca*), Eider (*Somateria*), Elegant tern (*Thalasseus elegans*), Emperor goose (*Anser canagicus*), Emu (*Dromaius novaehollandiae*), Eurasian collared dove (*Streptopelia decaocto*), Eurasian coot (*Fulica atra*), Eurasian curlew (*Numenius arquata*), Eurasian eagle-owl (*Bubo bubo*), Eurasian griffon vulture (*Gyps fulvus*), Eurasian hobby (*Falco subbuteo*), Eurasian jay (*Garrulus glandarius*), Eurasian oystercatcher (*Haematopus ostralegus*), Eurasian scops owl (*Otus scops*), Eurasian sparrowhawk (*Accipiter nisus*), Eurasian spoonbill (*Platalea leucorodia*), Eurasian wigeon (*Mareca penelope*), Eurasian woodcock (*Scolopax rusticola*), European polecat (*Mustela putorius*), European robin (*Erithacus rubecula*), European starling (*Sturnus vulgaris*), European turtle dove (*Streptopelia turtur*), Falcated duck (*Mareca falcata*), Falcon (*Falco*), Ferret (*Mustela furo*), Ferruginous hawk (*Buteo regalis*), Finch (Fringillidae), Fish crow (*Corvus ossifragus*), Fisher (*Pekania pennanti*), Fox (Canidae), Franklin's gull (*Leucophaeus pipixcan*), Frigatebird (*Fregata*), Fulmar (*Fulmarus*), Fur seal (*Arctocephalus australis*), Gadwall (*Mareca strepera*), Gannet (*Morus*), Garganey (*Spatula querquedula*), Geoffroy's cat (*Leopardus geoffroyi*), Glaucous gull (*Larus hyperboreus*), Glaucous-winged gull (*Larus glaucescens*), Goat (*Capra hircus*), Golden eagle (*Aquila chrysaetos*), Golden pheasant (*Chrysolophus pictus*), Goose (*Anser*), Grackle (Icteridae), Great black-backed gull (*Larus marinus*), Great black-hawk (*Buteogallus urubitinga*), Great black-headed gull (*Ichthyaetus ichthyaetus*), Great blue heron (*Ardea herodias*), Great cormorant (*Phalacrocorax carbo*), Great crested grebe (*Podiceps cristatus*), Great grey owl (*Strix nebulosa*), Great horned owl (*Bubo virginianus*), Great knot (*Calidris tenuirostris*), Great shearwater (*Ardenna gravis*), Great skua (*Stercorarius skua*), Great tit (*Parus major*), Great white pelican (*Pelecanus onocrotalus*), Great-tailed grackle (*Quiscalus mexicanus*), Greater flamingo (*Phoenicopterus roseus*), Greater rhea (*Rhea americana*), Greater scaup (*Aythya marila*), Greater yellowlegs (*Tringa melanoleuca*), Grebe (*Podiceps*), Green heron (*Butorides virescens*), Green pheasant (*Phasianus versicolor*), Green sandpiper (*Tringa ochropus*), Green-winged teal (*Anas carolinensis*), Grey goose (*Anser*), Grey gull (*Leucophaeus modestus*), Grey heron (*Ardea cinerea*), Grey seal (*Halichoerus grypus*), Grey teal (*Anas gracilis*), Grey-crowned crane (*Balearica regulorum*), Grey-faced buzzard (*Butastur indicus*), Grey-headed gull (*Chroicocephalus cirrocephalus*), Greylag goose (*Anser anser*), Guanay cormorant (*Leucocarbo bougainvillii*), Guinea (*Numida*), Guineafowl (Numididae), Gull (*Larus*), Gull-billed tern (*Gelochelidon nilotica*), Gyrfalcon (*Falco rusticolus*), Harbor porpoise (*Phocoena phocoena*), Harbor seal (*Phoca vitulina*), Harris's hawk (*Parabuteo unicinctus*), Hartlaub's gull (*Chroicocephalus hartlaubii*), Hawaiian goose (*Branta*), Hawk (Accipitridae), Helmeted guineafowl (*Numida meleagris*), Hen harrier (*Circus cyaneus*), Herons (*Ardea*), Herring gull (*Larus argentatus*), Hooded crane (*Grus monacha*), Hooded merganser (*Lophodytes cucullatus*), Hooded vulture (*Necrosyrtes monachus*), Horned grebe (*Podiceps auritus*), Horse (*Equus caballus*), Human (*Homo sapiens*), Humboldt penguin (*Spheniscus humboldti*), Ibis (*Threskiornis*), Iceland gull (*Larus glaucoides*), Inca tern (*Larosterna inca*), Indian peafowl (*Pavo cristatus*), Jack snipe (*Lymnocryptes minimus*), Jackal buzzard (*Buteo rufofuscus*), Jackdaw (*Coloeus monedula*), Japanese white-eye (*Zosterops japonicus*), Kalij pheasant (*Lophura leucomelanos*), Kelp gull (*Larus dominicanus*), King penguin (*Aptenodytes patagonicus*), King vulture (*Sarcoramphus papa*), Kittiwake (*Rissa*), Lady Amherst's pheasant (*Chrysolophus amherstiae*), Lapwing (*Vanellus*), Large-billed crow (*Corvus macrorhynchos*), Laughing gull (*Leucophaeus atricilla*), Least sandpiper (*Calidris minutilla*), Leopard (*Panthera pardus*), Leopard cat (*Prionailurus bengalensis*), Lesser black-backed gull (*Larus fuscus*), Lesser kestrel (*Falco naumanni*), Lesser scaup (*Aythya affinis*), Lesser white-fronted goose (*Anser erythropus*), Lion (*Panthera leo*), Little auk (*Alle alle*), Little egret (*Egretta garzetta*), Little grebe (*Tachybaptus ruficollis*), Little gull (*Hydrocoloeus minutus*), Little owl (*Athene noctua*), Little tern (*Sternula albifrons*), Long-eared owl (*Asio otus*), Long-tailed shrike (*Lanius schach*), Loon (*Gavia*), Lynx (*Lynx*), Macaque (*Macaca*), Magnificent frigatebird (*Fregata magnificens*), Magpie (Corvidae), Magpie robin (*Copsychus*), Mallard (*Anas platyrhynchos*), Mandarin duck (*Aix galericulata*), Manx shearwater (*Puffinus puffinus*), Marabou stork (*Leptoptilos crumenifer*), Marine otter (*Lontra felina*), Mediterranean gull (*Ichthyaetus melanocephalus*), Meerkat (*Suricata suricatta*), Merganser (*Mergus*), Merlin (*Falco columbarius*), Mexican duck (*Anas diazi*), Mink (*Neogale vison*), Moorhen (*Gallinula*), Mottled duck (*Anas fulvigula*), Mountain hawk-eagle (*Nisaetus nipalensis*), Mountain lion (*Puma concolor*), Mouse (*Mus musculus*), Munia (*Lonchura*), Muscovy duck (*Cairina moschata*), Mute swan (*Cygnus olor*), Nene (*Branta*), Neotropic cormorant (*Phalacrocorax brasilianus*), Night heron (*Nycticorax*), North american opossum (*Didelphis virginiana*), Northern fulmar (*Fulmarus glacialis*), Northern fur seal (*Callorhinus ursinus*), Northern gannet (*Morus bassanus*), Northern giant petrel (*Macronectes halli*), Northern goshawk (*Astur gentilis*), Northern harrier (*Circus hudsonius*), Northern lapwing (*Vanellus vanellus*), Northern pintail (*Anas acuta*), Northern shoveler (*Spatula clypeata*), Northwestern crow (*Corvus caurinus*), Norway rat (*Rattus norvegicus*), Oriental magpie-robin (*Copsychus saularis*), Osprey (*Pandion haliaetus*), Ostrich (*Struthio camelus*), Otter (*Lutra*), Owl (Strigidae), Owston's civet (*Chrotogale owstoni*), Oystercatcher (*Haematopus*), Pacific black duck (*Anas superciliosa*), Painted stork (*Mycteria leucocephala*), Pallas's sandgrouse (*Syrrhaptes paradoxus*), Parasitic jaeger (*Stercorarius parasiticus*), Parrot (Psittacidae), Partridge (*Perdicinae*), Pelican (Pelecanidae), Penguin (*Pygoscelis*), Peregrine falcon (*Falco peregrinus*), Peruvian booby (*Sula variegata*), Peruvian pelican (*Pelecanus thagus*), Pheasant (Phasianidae), Pigeon (Columbidae), Pigeon guillemot (*Cepphus columba*), Pika (*Ochotona*), Pin-tailed snipe (*Gallinago stenura*), Pine marten (*Martes martes*), Pink-footed goose (*Anser brachyrhynchus*), Pinniped (Pinnipedia), Pintado petrel (*Daption capense*), Plateau pika (*Ochotona curzoniae*), Polar bear (*Ursus maritimus*), Polecat (*Mustela*), Pomeranian goose (*Anser*), Pond heron (*Ardeola*), Porpoise (*Phocoena*), Purple heron (*Ardea purpurea*), Quail (Phasianidae), Raccoon (*Procyon lotor*), Raccoon dog (*Nyctereutes procyonoides*), Raptor (Telluraves), Raven (*Corvus*), Razorbill (*Alca torda*), Red fox (*Vulpes vulpes*), Red kite (*Milvus milvus*), Red knot (*Calidris canutus*), Red-backed hawk (*Geranoaetus polyosoma*), Red-billed blue magpie (*Urocissa erythroryncha*), Red-billed starling (*Spodiopsar sericeus*), Red-breasted goose (*Branta ruficollis*), Red-breasted merganser (*Mergus serrator*), Red-crested pochard (*Netta rufina*), Red-crowned crane (*Grus japonensis*), Red-necked grebe (*Podiceps grisegena*), Red-necked phalarope (*Tringa tobata*), Red-shouldered hawk (*Buteo lineatus*), Red-tailed hawk (*Buteo jamaicensis*), Redhead duck (*Aythya americana*), Reed warbler (*Acrocephalus*), Rhea (*Rhea*), Rhinoceros auklet (*Cerorhinca monocerata*), Ring-billed gull (*Larus delawarensis*), Ring-necked duck (*Aythya collaris*), Ringed seal (*Pusa hispida*), Rock pigeon (*Columba livia*), Rook (*Corvus frugilegus*), Roseate spoonbill (*Platalea ajaja*), Ross's goose (*Anser rossii*), Rosy-billed pochard (*Netta peposaca*), Rough-legged hawk (*Buteo lagopus*), Royal tern (*Thalasseus maximus*), Ruddy shelduck (*Tadorna ferruginea*), Ruddy turnstone (*Arenaria interpres*), Ruffed grouse (*Bonasa umbellus*), Rufous hornero (*Furnarius rufus*), Sabine's gull (*Xema sabini*), Sacred ibis (*Threskiornis aethiopicus*), Saker falcon (*Falco cherrug*), Sanderling (*Calidris alba*), Sandhill crane (*Antigone canadensis*), Sandpiper (Scolopacidae), Sandwich tern (*Thalasseus sandvicensis*), Scaly-breasted munia (*Lonchura punctulata*), Scarlet ibis (*Eudocimus ruber*), Scarlet macaw (*Ara macao*), Scoter (*Melanitta*), Sea eagle (*Haliaeetus*), Sea lion (*Zalophus*), Sea otter (*Enhydra lutris*), Seabird (Aves), Seal (*Phoca*), Serval (*Leptailurus serval*), Shag (*Phalacrocorax aristotelis*), Sharp-shinned hawk (*Accipiter striatus*), Shearwater (*Ardenna*), Shelduck (*Tadorna*), Shorebird (Charadriiformes), Short-billed gull (*Larus brachyrhynchus*), Short-eared owl (*Asio flammeus*), Short-tailed shearwater (*Ardenna tenuirostris*), Shrike (*Lanius*), Siberian blue robin (*Emberiza variabilis*), Silver pheasant (*Lophura nycthemera*), Silver teal (*Spatula versicolor*), Silver-eared mesia (*Leiothrix argentauris*), Skua (*Stercorarius*), Skunk (Mephitidae), Slaty-backed gull (*Larus schistisagus*), Slender-billed gull (*Chroicocephalus genei*), Smew (*Mergellus albellus*), Snow goose (*Anser caerulescens*), Snowy egret (*Egretta thula*), Snowy owl (*Bubo scandiacus*), Snowy plover (*Anarhynchus nivosus*), Song thrush (*Turdus philomelos*), South American coati (*Nasua nasua*), South American sea lion (*Otaria byronia*), South American tern (*Sterna hirundinacea*), South Georgia shag (*Leucocarbo georgianus*), South polar skua (*Stercorarius maccormicki*), Southern elephant seal (*Mirounga leonina*), Southern fulmar (*Fulmarus glacialoides*), Sparrow (Passeridae), Speckled pigeon (*Columba guinea*), Spectacled guillemot (*Cepphus carbo*), Spoonbill (*Platalea*), Spot-billed duck (*Anas poecilorhyncha*), Spotted eagle owl (*Bubo africanus*), Spur-winged goose (*Plectropterus gambensis*), Square-tailed nightjar (*Caprimulgus fossii*), Steamer duck (*Tachyeres*), Stone curlew (*Burhinus*), Stone marten (*Martes foina*), Stork (*Ciconia*), Striped skunk (*Mephitis mephitis*), Surf scoter (*Melanitta perspicillata*), Swainson's hawk (*Buteo swainsoni*), Swan (*Cygnus*), Swan goose (*Anser cygnoides*), Swift tern (*Thalasseus bergii*), Swiftlet (*Aerodramus*), Swine (*Sus scrofa*), Tawny owl (*Strix aluco*), Teal (*Anas*), Tern (Laridae), Thayer's gull (*Larus thayeri*), Thick-billed murre (*Uria lomvia*), Thrush (Turdidae), Tibetan blue bear (*Ursus arctos*), Tiger (*Panthera tigris*), Tree sparrow (*Passer montanus*), Tropical screech owl (*Megascops choliba*), Tropical shearwater (*Puffinus bailloni*), Trumpeter swan (*Cygnus buccinator*), Tufted duck (*Aythya fuligula*), Tundra bean goose (*Anser serrirostris*), Tundra swan (*Cygnus columbianus*), Turkey (*Meleagris gallopavo*), Turkey vulture (*Cathartes aura*), Turtle dove (*Streptopelia*), Unknown bird (Aves), Ural owl (*Strix uralensis*), Von schrenck's bittern (*Ixobrychus eurhythmus*), Waterbird (Aves), Watercock (*Gallicrex cinerea*), Waterfowl (Anatidae), Western barn owl (*Tyto alba*), Western cattle egret (*Bubulcus ibis*), Western grebe (*Aechmophorus occidentalis*), Western gull (*Larus occidentalis*), Western marsh harrier (*Circus aeruginosus*), Western sandpiper (*Calidris mauri*), Western screech owl (*Megascops kennicottii*), Whiskered tern (*Chlidonias hybrida*), Whistling duck (*Dendrocygna*), White mink (*Mustela erminea*), White stork (*Ciconia ciconia*), White-backed munia (*Lonchura striata*), White-backed stilt (*Himantopus himantopus*), White-breasted cormorant (*Phalacrocorax lucidus*), White-eyed pochard (*Aythya nyroca*), White-faced ibis (*Plegadis chihi*), White-faced whistling duck (*Dendrocygna viduata*), White-fronted goose (*Anser albifrons*), White-naped crane (*Antigone vipio*), White-tailed eagle (*Haliaeetus albicilla*), White-winged black tern (*Chlidonias leucopterus*), White-winged dove (*Zenaida asiatica*), White-winged scoter (*Melanitta fusca*), Whooper swan (*Cygnus cygnus*), Whooping crane (*Grus americana*), Willet (*Tringa semipalmata*), Wood duck (*Aix sponsa*), Yellow-billed duck (*Anas undulata*), Yellow-billed pintail (*Anas georgica*), Yellow-billed teal (*Anas flavirostris*), Yellow-headed amazon (*Amazona oratrix*), Yellow-legged gull (*Larus michahellis*)

**Table S4.** Countries with data in global regions within the cleaned and de-duplicated dataset. Regions were defined by R packages ‘countrycode’ based on the UN M49 geoscheme.

| **Global region** | **Countries** |
| --- | --- |
| Western Europe | Austria, Belgium, France, Germany, Luxembourg, Netherlands, Switzerland |
| Northern Europe | Denmark, Estonia, Finland, Iceland, Ireland, Isle of Man, Jersey, Latvia, Lithuania, Norway, Svalbard and Jan Mayen, Sweden, United Kingdom |
| Southern Europe | Albania, Bosnia and Herzegovina, Croatia, Greece, Italy, Kosovo, Macedonia, Montenegro, Portugal, Serbia, Slovenia, Spain |
| Eastern Europe | Bulgaria, Czech Republic, Hungary, Moldova, Poland, Romania, Russia, Slovakia, Ukraine |
| Western Africa | Benin, Burkina Faso, Cote d’Ivoire, Gambia, Ghana, Mali, Mauritania, Niger, Nigeria, Senegal, Togo |
| Northern Africa | Algeria, Egypt, Libya, Sudan |
| Southern Africa | Botswana, Lesotho, Namibia, South Africa |
| Eastern Africa | Madagascar, Mozambique, Somalia, Uganda, Zimbabwe |
| Middle Africa | Cameroon, Democratic Republic of the Congo, Gabon |
| Western Asia | Azerbaijan, Cyprus, Georgia, Iraq, Israel, Kuwait, Lebanon, Palestine, Saudi Arabia, Turkey, United Arab Emirates, Yemen |
| Central Asia | Kazakhstan |
| Southern Asia | Afghanistan, Bangladesh, Bhutan, India, Iran, Maldives, Nepal, Pakistan, Sri Lanka |
| South-Eastern Asia | Cambodia, Indonesia, Laos, Malaysia, Myanmar, Philippines, Singapore, Thailand, Vietnam |
| Eastern Asia | China, Hong Kong, Japan, Mongolia, North Korea, South Korea, Taiwan |
| North America | Canada, Greenland, United States |
| Central America | Costa Rica, El Salvador, Guatemala, Honduras, Mexico, Panama |
| Caribbean | Cayman Islands, Dominican Republic, Puerto Rico |
| South America | Argentina, Bolivia, Brazil, Chile, Colombia, Ecuador, Falkland Islands, Peru, Uruguay, Venezuela |
| Oceania | Australia, New Zealand |
| Antarctica | Antarctica, French Southern Territories, Livingston Island, Roberts Island, South Georgia and the South Sandwich Islands, Torgersen Island |

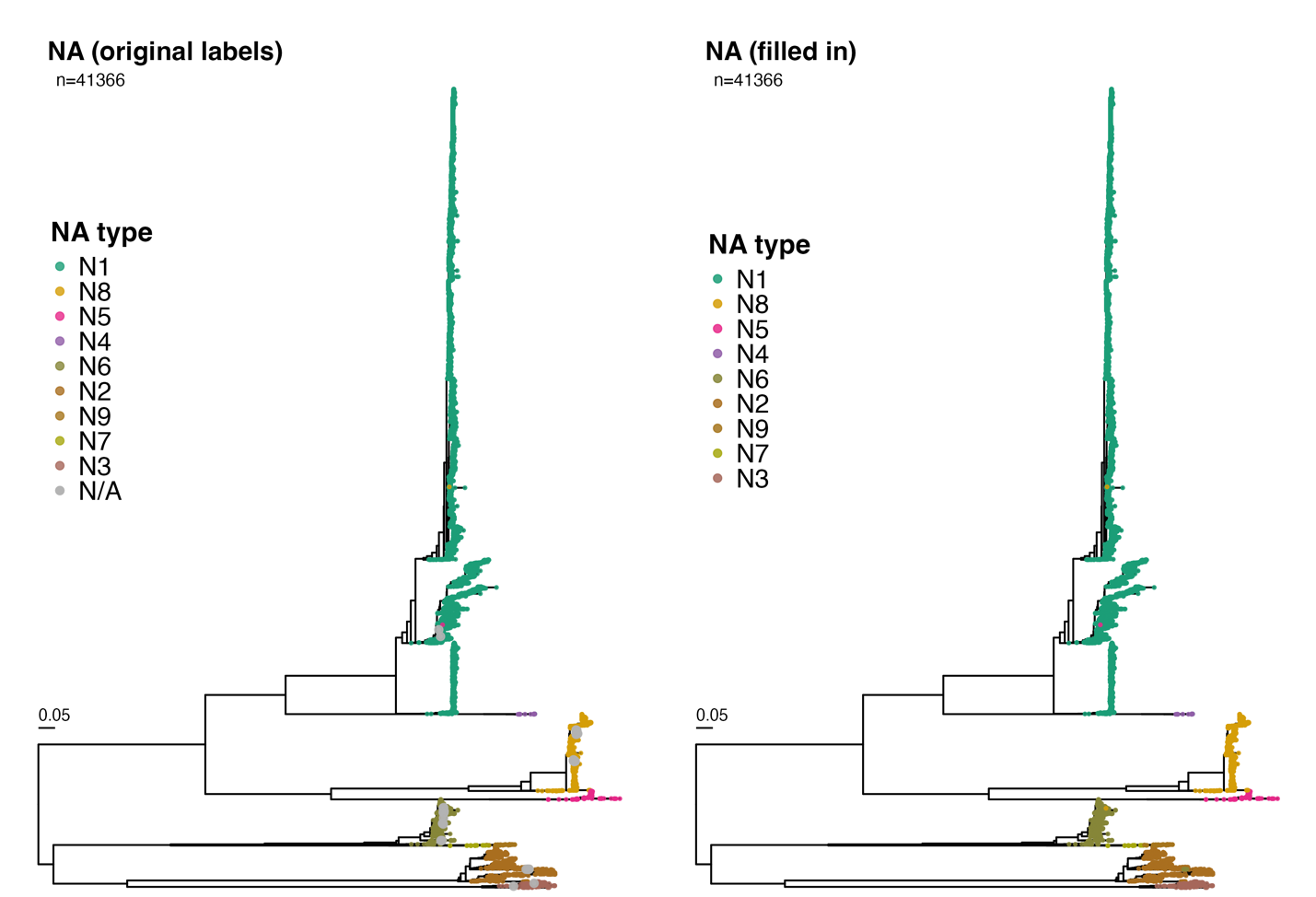

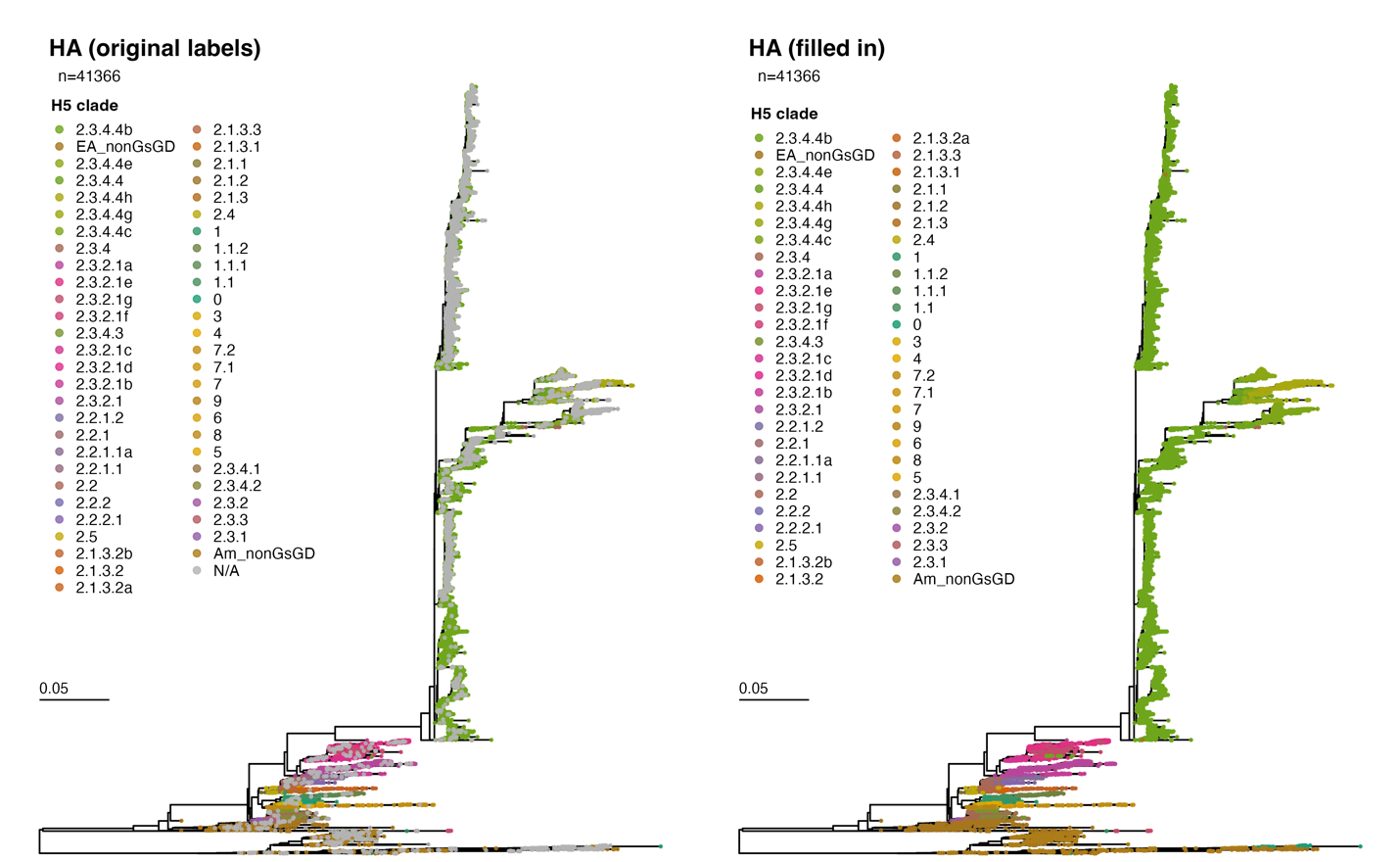
**Fig. S4**. Maximum likelihood HA phylogeny colored by H5 clade before and after filling in labels based on a phylogenetic neighbourhood algorithm. Clades in legend were sorted by tree order.

**Fig. S5**. Maximum likelihood NA phylogeny colored by NA type before and after filling in labels using a phylogenetic neighbourhood algorithm. Types in legend were sorted by tree order.

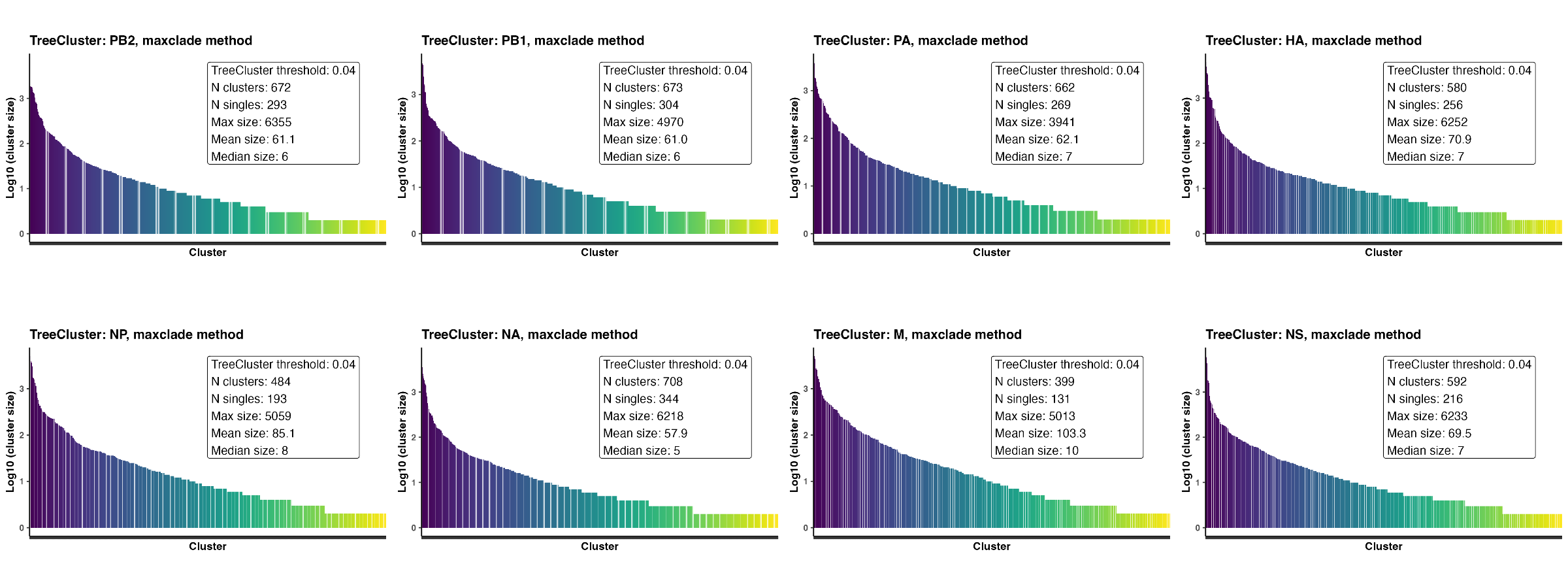

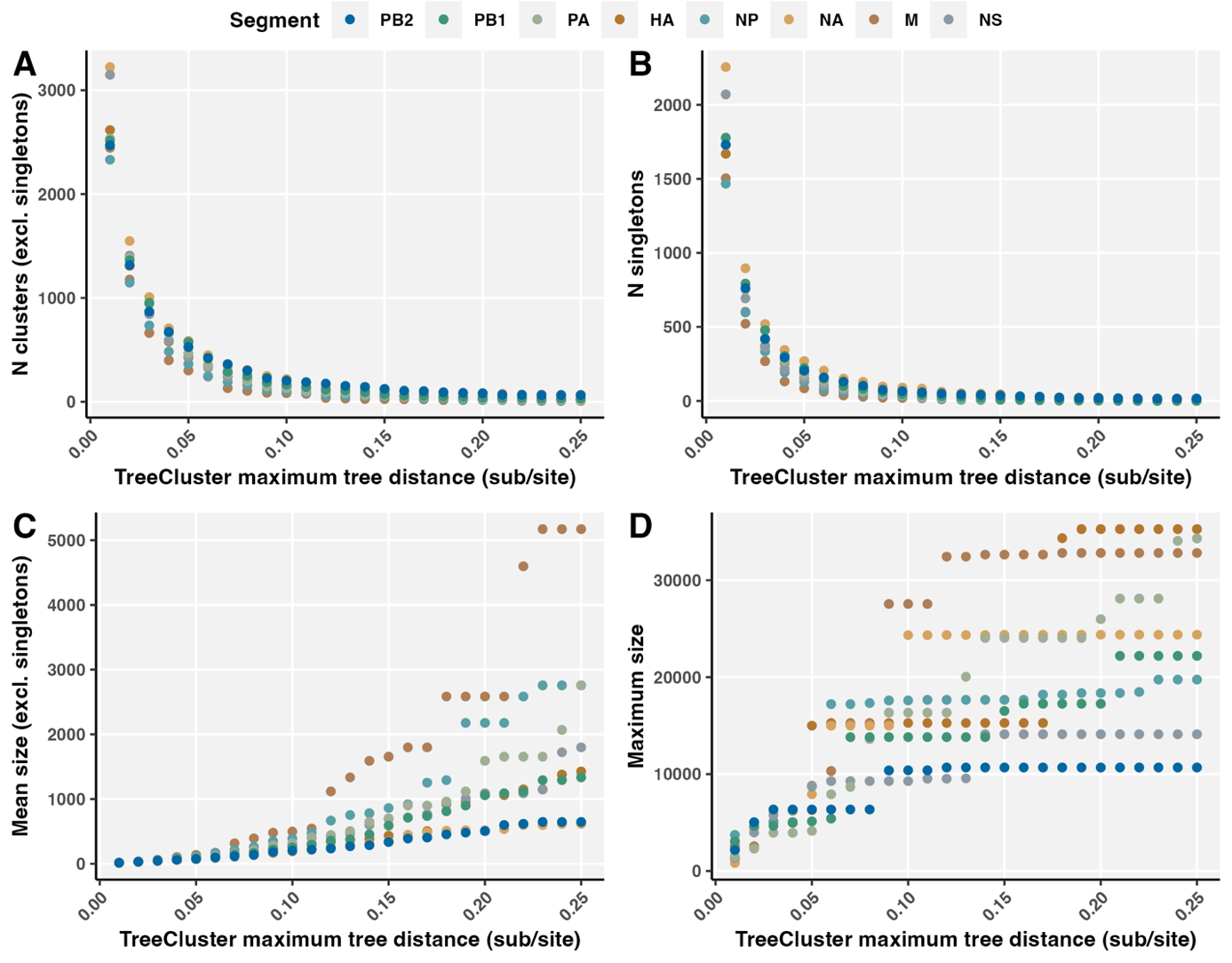
 **Fig. S6.** Evaluation of TreeCluster maximum pairwise patristic distance across segments. The A) number of clusters (excluding singletons), B) number of singletons, C) mean cluster size (excluding singletons), and D) maximum cluster size, for maximum patristic tree distances 0.01 to 0.25 substitutions/site (subs/site).

**Fig. S7.** Phylogenetic cluster size distribution with TreeCluster threshold: 0.04 subs/site. Annotated with number of clusters and singletons, and the maximum, mean, and median size excluding singletons.

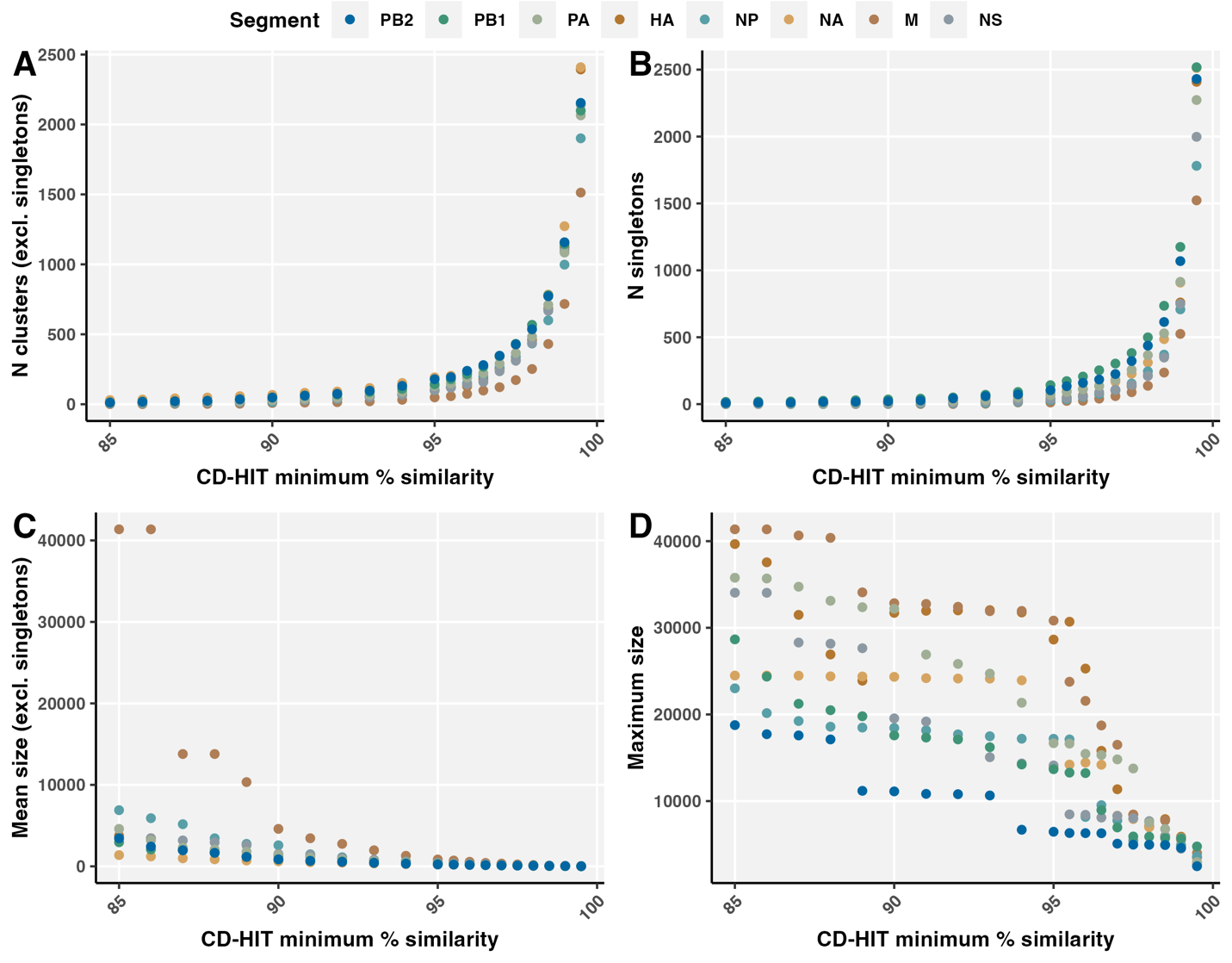
**Fig. S8.** Evaluation of CD-HIT clusters at 85-99% minimum genetic similarity across segments. The A) number of clusters (excluding singletons), B) number of singletons, C) mean cluster size (excluding singletons), and D) maximum cluster size, for genetic similarity thresholds 85-99.5%.

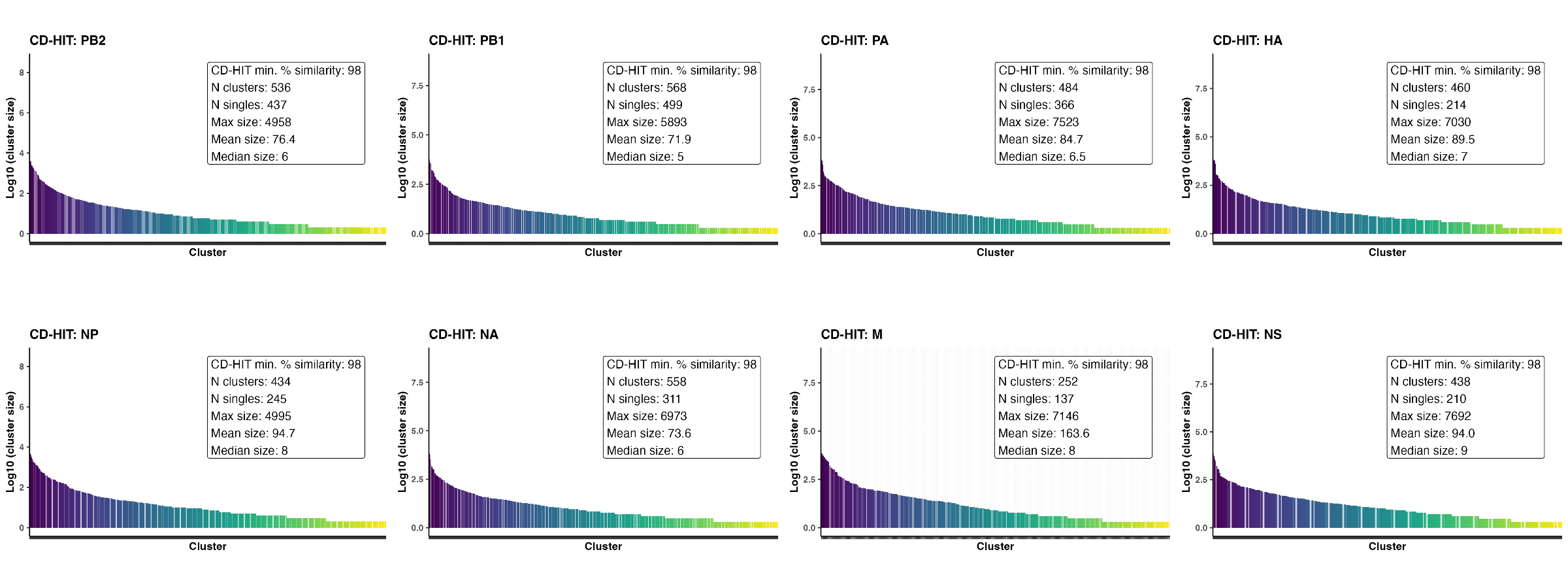

**Fig. S9.** Genetic cluster size distribution from CD-HIT with 98% similarity threshold. Annotated with number of clusters and singletons, and the maximum, mean, and median size excluding singletons.

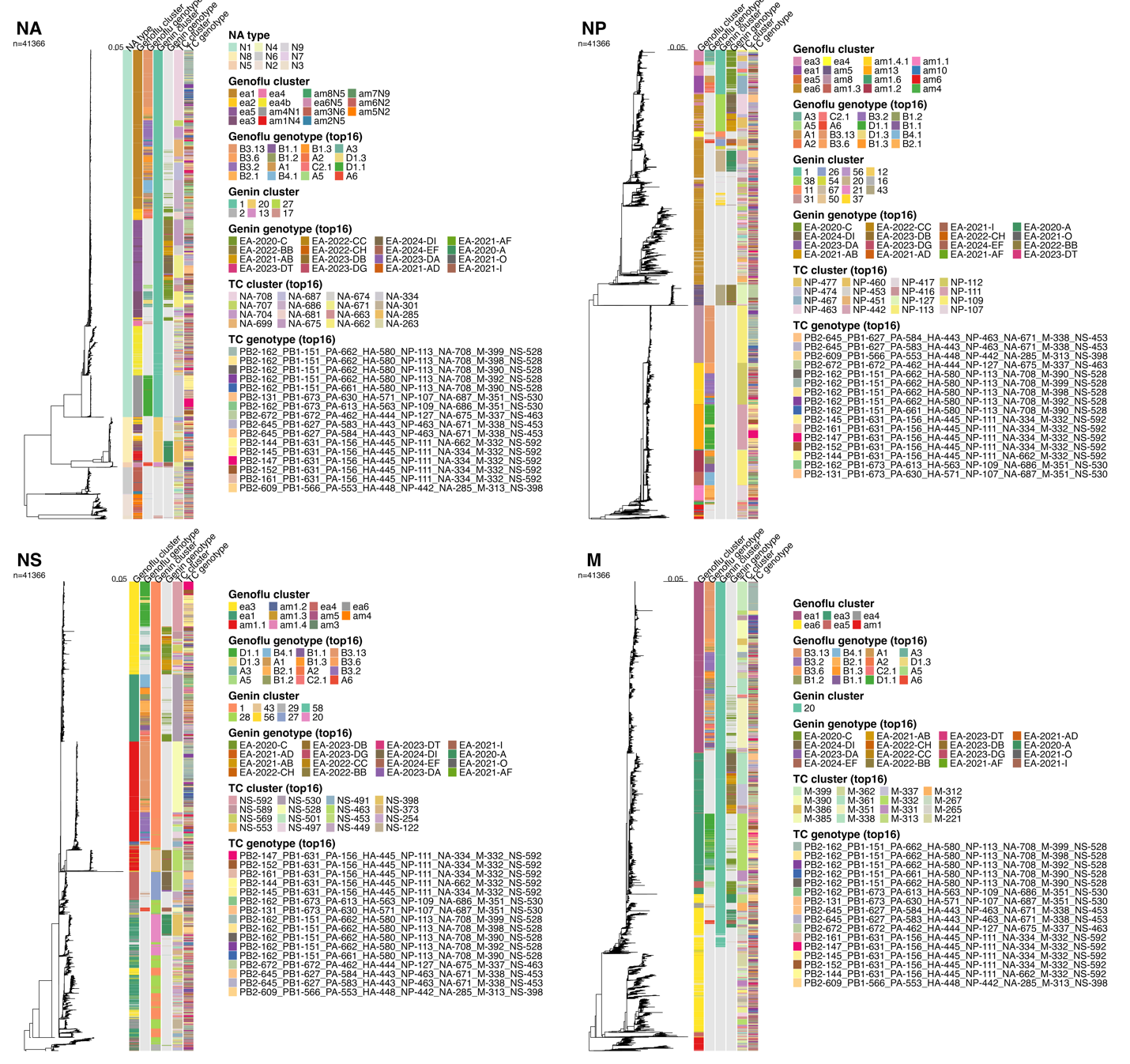

**Fig. S10.** Maximum likelihood phylogenies for segments NA, NP, M, and NS (n=41,366) comparing genotype and cluster labels for Genoflu, Genin2, and TreeCluster (TC). The NA tree also displays NA type. Top 16 most frequent labels were included in legends. Segments PB2, PB1, PA, and HA in Fig. 4.

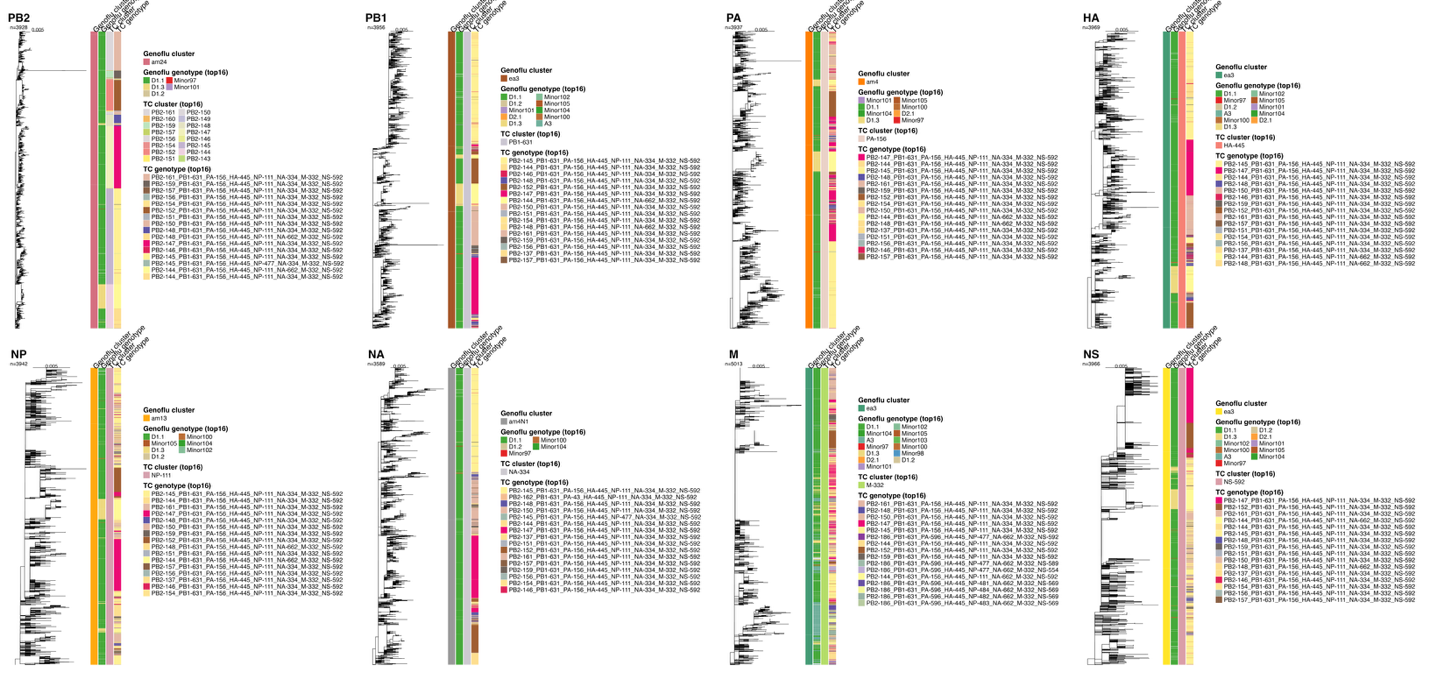
**Fig. S11.** Maximum likelihood trees for GenoFlu genotype D1.1. Subtrees clipped at most recent common ancestor sharing genotype label. Tile annotations for GenoFlu segment cluster and genotype, and TreeCluster segment cluster and genotypes. Top 16 most frequent labels were included in legends.

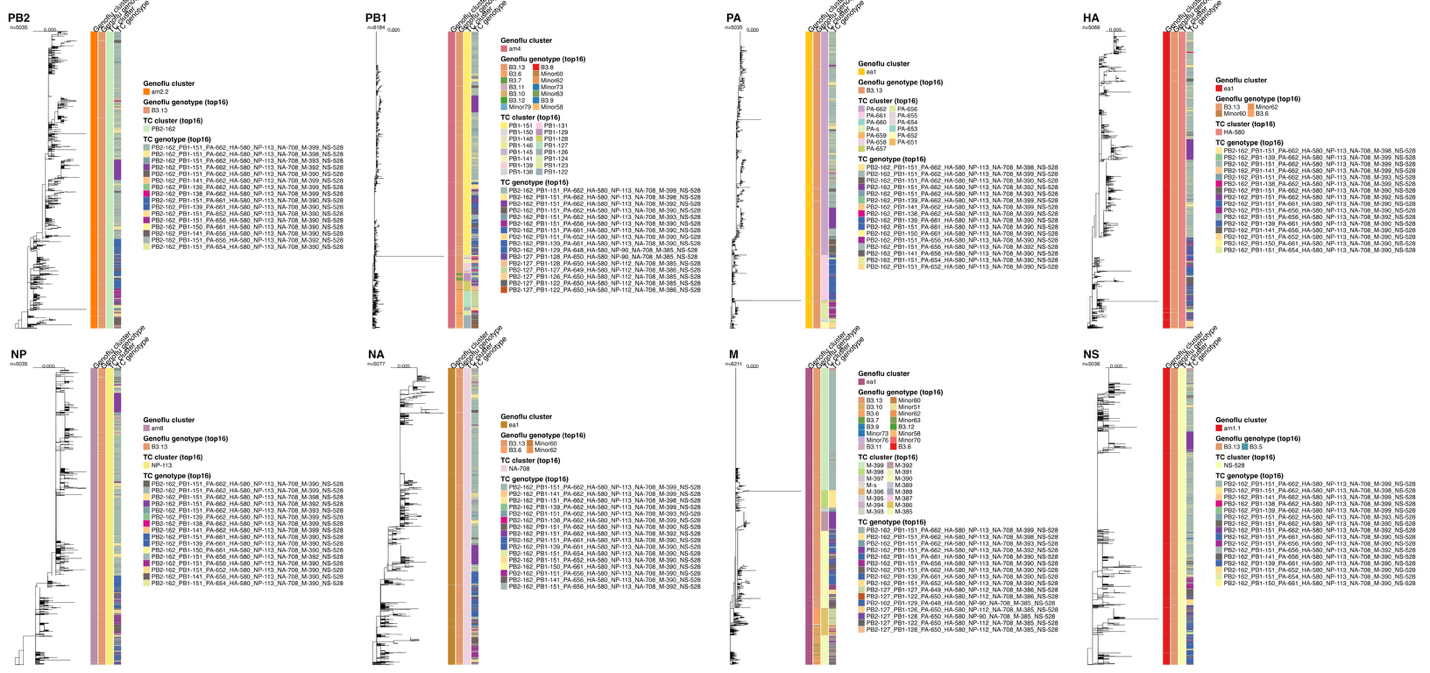

**Fig. S12.** Maximum likelihood trees for GenoFlu genotype B3.13. Subtrees clipped at most recent common ancestor sharing genotype label. Tile annotations for GenoFlu segment cluster and genotype, and TreeCluster segment cluster and genotypes. Top 16 most frequent labels were included in legends.

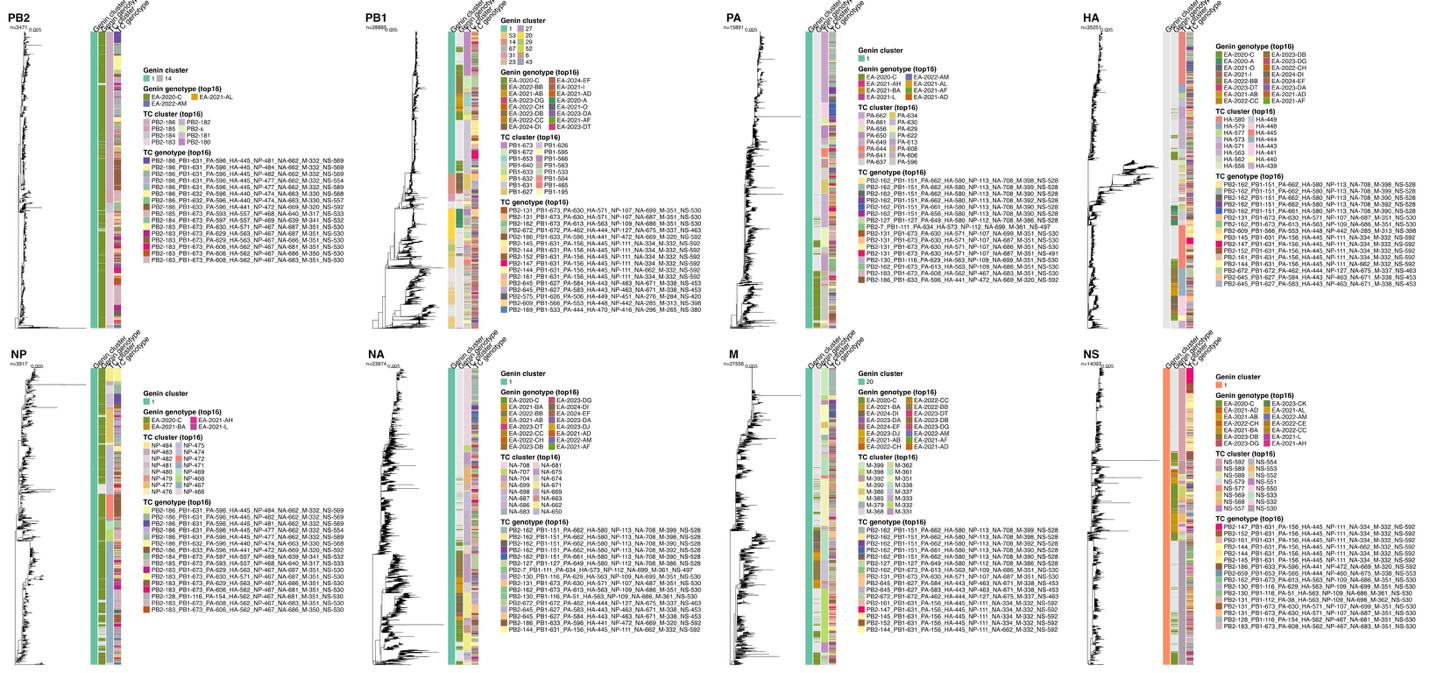

**Fig. S13.** Maximum likelihood trees for Genin2 genotype EA-2020-C. Subtrees clipped at most recent common ancestor sharing genotype label. Tile annotations for Genin2 segment cluster and genotype, and TreeCluster segment cluster and genotypes. Top 16 most frequent labels were included in legends. Note that Genin2 does not assign HA clusters.

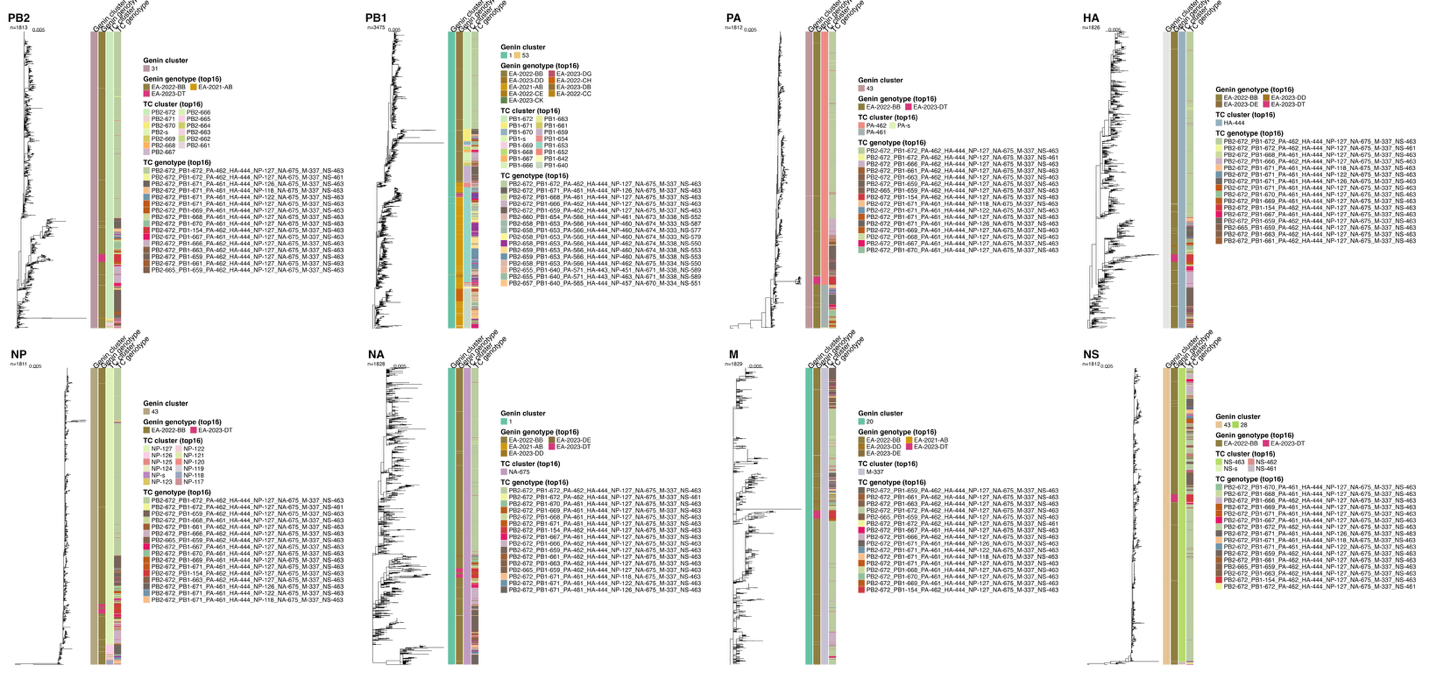

**Fig. S14.** Maximum likelihood trees for Genin2 genotype EA-2022-BB. Subtrees clipped at most recent common ancestor sharing genotype label. Tile annotations for Genin2 segment cluster and genotype, and TreeCluster segment cluster and genotypes. Top 16 most frequent labels were included in legends. Note that Genin2 does not assign HA clusters.

**Table S5.** Phylogenetic cluster composition of GenoFlu genotypes with at least 10 sequences. Clusters were numbered from bottom to top of trees. Number of sequences and number of TreeCluster (TC) genotypes; for each segment, the number (N) of TreeCluster clusters, cluster IDs range (minimum to maximum cluster ID), cluster ID span (maximum ID – minimum ID + 1), and the span-to-cluster ratio. Number and range of clusters exclude singletons; number of genotypes (genos) includes singletons.

|  |  |  | **PB2** | | | | **PB1** | | | | **PA** | | | | **HA** | | | | **NP** | | | | **NA** | | | | **M** | | | | **NS** | | | |
| --- | --- | --- | --- | --- | --- | --- | --- | --- | --- | --- | --- | --- | --- | --- | --- | --- | --- | --- | --- | --- | --- | --- | --- | --- | --- | --- | --- | --- | --- | --- | --- | --- | --- | --- |
| **GenoFlu**  **genotype** | N seq | N TC geno | N clust | ID range | Span | Span  /N | N clust | ID range | Span | Span  /N | N clust | ID range | Span | Span  /N | N clust | ID range | Span | Span  /N | N clust | ID range | Span | Span  /N | N clust | ID range | Span | Span  /N | N clust | ID range | Span | Span  /N | N clust | ID range | Span | Span  /N |
| **B3.13** | 4985 | 85 | 1 | PB2-162 | 1 | 1 | 22 | PB1-122–PB1-151 | 30 | 1.4 | 12 | PA-651–PA-662 | 12 | 1 | 1 | HA-580 | 1 | 1 | 1 | NP-113 | 1 | 1 | 1 | NA-708 | 1 | 1 | 12 | M-385–M-399 | 15 | 1.2 | 1 | NS-528 | 1 | 1 |
| **D1.1** | 3525 | 21 | 20 | PB2-142–PB2-161 | 20 | 1 | 1 | PB1-631 | 1 | 1 | 1 | PA-156 | 1 | 1 | 1 | HA-445 | 1 | 1 | 1 | NP-111 | 1 | 1 | 1 | NA-334 | 1 | 1 | 1 | M-332 | 1 | 1 | 1 | NS-592 | 1 | 1 |
| **B3.2** | 2173 | 244 | 15 | PB2-2–PB2-20 | 19 | 1.3 | 1 | PB1-111 | 1 | 1 | 21 | PA-611–PA-645 | 35 | 1.7 | 10 | HA-563–HA-580 | 18 | 1.8 | 1 | NP-112 | 1 | 1 | 12 | NA-686–NA-708 | 23 | 1.9 | 23 | M-350–M-398 | 49 | 2.1 | 33 | NS-496–NS-528 | 33 | 1 |
| **B4.1** | 1121 | 36 | 1 | PB2-162 | 1 | 1 | 1 | PB1-673 | 1 | 1 | 16 | PA-612–PA-632 | 21 | 1.3 | 1 | HA-563 | 1 | 1 | 1 | NP-109 | 1 | 1 | 3 | NA-686–NA-699 | 14 | 4.7 | 11 | M-348–M-367 | 20 | 1.8 | 1 | NS-530 | 1 | 1 |
| **B2.1** | 1014 | 58 | 1 | PB2-131 | 1 | 1 | 1 | PB1-673 | 1 | 1 | 6 | PA-619–PA-630 | 12 | 2 | 10 | HA-563–HA-573 | 11 | 1.1 | 1 | NP-107 | 1 | 1 | 8 | NA-684–NA-702 | 19 | 2.4 | 11 | M-347–M-367 | 21 | 1.9 | 1 | NS-530 | 1 | 1 |
| **A3** | 971 | 27 | 1 | PB2-186 | 1 | 1 | 2 | PB1-630–PB1-631 | 2 | 1 | 2 | PA-595–PA-596 | 2 | 1 | 2 | HA-439–HA-445 | 7 | 3.5 | 11 | NP-474–NP-484 | 11 | 1 | 2 | NA-645–NA-662 | 18 | 9 | 2 | M-331–M-332 | 2 | 1 | 11 | NS-551–NS-592 | 42 | 3.8 |
| **B3.6** | 884 | 42 | 1 | PB2-127 | 1 | 1 | 12 | PB1-119–PB1-130 | 12 | 1 | 5 | PA-646–PA-650 | 5 | 1 | 1 | HA-580 | 1 | 1 | 1 | NP-112 | 1 | 1 | 2 | NA-707–NA-708 | 2 | 1 | 6 | M-381–M-388 | 8 | 1.3 | 4 | NS-513–NS-528 | 16 | 4 |
| **B1.1** | 699 | 18 | 1 | PB2-130 | 1 | 1 | 1 | PB1-116 | 1 | 1 | 2 | PA-629–PA-630 | 2 | 1 | 2 | HA-563–HA-571 | 9 | 4.5 | 1 | NP-109 | 1 | 1 | 7 | NA-684–NA-699 | 16 | 2.3 | 5 | M-350–M-361 | 12 | 2.4 | 1 | NS-530 | 1 | 1 |
| **B1.2** | 646 | 39 | 1 | PB2-130 | 1 | 1 | 1 | PB1-116 | 1 | 1 | 1 | PA-51 | 1 | 1 | 2 | HA-563–HA-571 | 9 | 4.5 | 1 | NP-109 | 1 | 1 | 5 | NA-686–NA-699 | 14 | 2.8 | 18 | M-350–M-369 | 20 | 1.1 | 1 | NS-530 | 1 | 1 |
| **A1** | 581 | 49 | 1 | PB2-183 | 1 | 1 | 1 | PB1-673 | 1 | 1 | 11 | PA-610–PA-631 | 22 | 2 | 11 | HA-563–HA-573 | 11 | 1 | 1 | NP-467 | 1 | 1 | 6 | NA-684–NA-699 | 16 | 2.7 | 7 | M-350–M-361 | 12 | 1.7 | 1 | NS-530 | 1 | 1 |
| **B1.3** | 529 | 10 | 1 | PB2-131 | 1 | 1 | 1 | PB1-112 | 1 | 1 | 1 | PA-38 | 1 | 1 | 1 | HA-563 | 1 | 1 | 1 | NP-109 | 1 | 1 | 9 | NA-687–NA-699 | 13 | 1.4 | 3 | M-351–M-362 | 12 | 4 | 1 | NS-530 | 1 | 1 |
| **A2** | 515 | 26 | 1 | PB2-183 | 1 | 1 | 1 | PB1-673 | 1 | 1 | 10 | PA-603–PA-629 | 27 | 2.7 | 2 | HA-561–HA-562 | 2 | 1 | 1 | NP-467 | 1 | 1 | 5 | NA-679–NA-686 | 8 | 1.6 | 6 | M-348–M-353 | 6 | 1 | 1 | NS-530 | 1 | 1 |
| **D1.3** | 352 | 3 | 3 | PB2-143–PB2-148 | 6 | 2 | 1 | PB1-631 | 1 | 1 | 1 | PA-156 | 1 | 1 | 1 | HA-445 | 1 | 1 | 1 | NP-111 | 1 | 1 | 1 | NA-662 | 1 | 1 | 1 | M-332 | 1 | 1 | 1 | NS-592 | 1 | 1 |
| **A6** | 315 | 1 | 1 | PB2-575 | 1 | 1 | 1 | PB1-626 | 1 | 1 | 1 | PA-506 | 1 | 1 | 1 | HA-449 | 1 | 1 | 1 | NP-451 | 1 | 1 | 1 | NA-276 | 1 | 1 | 1 | M-284 | 1 | 1 | 1 | NS-420 | 1 | 1 |
| **A5** | 285 | 42 | 1 | PB2-186 | 1 | 1 | 1 | PB1-632 | 1 | 1 | 1 | PA-596 | 1 | 1 | 1 | HA-440 | 1 | 1 | 1 | NP-474 | 1 | 1 | 1 | NA-663 | 1 | 1 | 7 | M-324–M-330 | 7 | 1 | 15 | NS-553–NS-569 | 17 | 1.1 |
| **C2.1** | 281 | 1 | 1 | PB2-128 | 1 | 1 | 1 | PB1-116 | 1 | 1 | 1 | PA-154 | 1 | 1 | 1 | HA-562 | 1 | 1 | 1 | NP-467 | 1 | 1 | 1 | NA-681 | 1 | 1 | 1 | M-351 | 1 | 1 | 1 | NS-530 | 1 | 1 |
| **B2.2** | 252 | 6 | 1 | PB2-131 | 1 | 1 | 1 | PB1-673 | 1 | 1 | 1 | PA-630 | 1 | 1 | 2 | HA-571–HA-573 | 3 | 1.5 | 1 | NP-107 | 1 | 1 | 4 | NA-686–NA-702 | 17 | 4.2 | 2 | M-348–M-351 | 4 | 2 | 1 | NS-491 | 1 | 1 |
| **B3.1** | 225 | 48 | 1 | PB2-7 | 1 | 1 | 1 | PB1-673 | 1 | 1 | 12 | PA-612–PA-634 | 23 | 1.9 | 4 | HA-563–HA-575 | 13 | 3.2 | 1 | NP-112 | 1 | 1 | 3 | NA-686–NA-699 | 14 | 4.7 | 8 | M-351–M-365 | 15 | 1.9 | 1 | NS-530 | 1 | 1 |
| **B3.5** | 153 | 4 | 1 | PB2-137 | 1 | 1 | 1 | PB1-111 | 1 | 1 | 2 | PA-634–PA-643 | 10 | 5 | 2 | HA-573–HA-579 | 7 | 3.5 | 1 | NP-112 | 1 | 1 | 2 | NA-699–NA-707 | 9 | 4.5 | 3 | M-361–M-380 | 20 | 6.7 | 3 | NS-497–NS-528 | 32 | 10.7 |
| **A4** | 144 | 12 | 6 | PB2-580–PB2-586 | 7 | 1.2 | 1 | PB1-503 | 1 | 1 | 1 | PA-586 | 1 | 1 | 2 | HA-456–HA-457 | 2 | 1 | 1 | NP-453 | 1 | 1 | 5 | NA-632–NA-637 | 6 | 1.2 | 1 | M-312 | 1 | 1 | 1 | NS-449 | 1 | 1 |
| **B3.7** | 110 | 3 | 1 | PB2-162 | 1 | 1 | 1 | PB1-129 | 1 | 1 | 1 | PA-648 | 1 | 1 | 1 | HA-580 | 1 | 1 | 1 | NP-90 | 1 | 1 | 1 | NA-708 | 1 | 1 | 3 | M-385–M-387 | 3 | 1 | 1 | NS-528 | 1 | 1 |
| **B3.10** | 108 | 5 | 1 | PB2-127 | 1 | 1 | 1 | PB1-128 | 1 | 1 | 1 | PA-650 | 1 | 1 | 1 | HA-580 | 1 | 1 | 1 | NP-90 | 1 | 1 | 1 | NA-708 | 1 | 1 | 4 | M-383–M-388 | 6 | 1.5 | 2 | NS-521–NS-528 | 8 | 4 |
| **B3.3** | 83 | 1 | 1 | PB2-140 | 1 | 1 | 1 | PB1-90 | 1 | 1 | 1 | PA-634 | 1 | 1 | 1 | HA-573 | 1 | 1 | 1 | NP-112 | 1 | 1 | 1 | NA-699 | 1 | 1 | 1 | M-351 | 1 | 1 | 1 | NS-497 | 1 | 1 |
| **B5.1** | 23 | 4 | 1 | PB2-183 | 1 | 1 | 1 | PB1-673 | 1 | 1 | 1 | PA-630 | 1 | 1 | 2 | HA-563–HA-571 | 9 | 4.5 | 1 | NP-112 | 1 | 1 | 2 | NA-686–NA-699 | 14 | 7 | 2 | M-351–M-353 | 3 | 1.5 | 1 | NS-530 | 1 | 1 |
| **B3.4** | 20 | 6 | 1 | PB2-95 | 1 | 1 | 1 | PB1-111 | 1 | 1 | 2 | PA-644–PA-645 | 2 | 1 | 2 | HA-579–HA-580 | 2 | 1 | 1 | NP-112 | 1 | 1 | 1 | NA-707 | 1 | 1 | 1 | M-380 | 1 | 1 | 5 | NS-512–NS-528 | 17 | 3.4 |
| **Minor100** | 13 | 1 | 1 | PB2-137 | 1 | 1 | 1 | PB1-631 | 1 | 1 | 1 | PA-156 | 1 | 1 | 1 | HA-445 | 1 | 1 | 1 | NP-111 | 1 | 1 | 1 | NA-334 | 1 | 1 | 1 | M-332 | 1 | 1 | 1 | NS-592 | 1 | 1 |
| **Minor62** | 13 | 2 | 1 | PB2-126 | 1 | 1 | 2 | PB1-122–PB1-123 | 2 | 1 | 1 | PA-650 | 1 | 1 | 1 | HA-580 | 1 | 1 | 1 | NP-112 | 1 | 1 | 1 | NA-708 | 1 | 1 | 2 | M-385–M-386 | 2 | 1 | 1 | NS-528 | 1 | 1 |
| **Minor87** | 13 | 1 | 1 | PB2-128 | 1 | 1 | 1 | PB1-116 | 1 | 1 | 1 | PA-154 | 1 | 1 | 1 | HA-562 | 1 | 1 | 1 | NP-107 | 1 | 1 | 1 | NA-681 | 1 | 1 | 1 | M-351 | 1 | 1 | 1 | NS-530 | 1 | 1 |
| **Minor81** | 12 | 1 | 1 | PB2-128 | 1 | 1 | 1 | PB1-110 | 1 | 1 | 1 | PA-606 | 1 | 1 | 1 | HA-562 | 1 | 1 | 1 | NP-97 | 1 | 1 | 1 | NA-681 | 1 | 1 | 1 | M-351 | 1 | 1 | 1 | NS-530 | 1 | 1 |
| **B3.12** | 10 | 1 | 1 | PB2-127 | 1 | 1 | 1 | PB1-128 | 1 | 1 | 1 | PA-650 | 1 | 1 | 1 | HA-580 | 1 | 1 | 1 | NP-111 | 1 | 1 | 1 | NA-708 | 1 | 1 | 1 | M-385 | 1 | 1 | 1 | NS-528 | 1 | 1 |
| **Minor19** | 10 | 1 | 1 | PB2-131 | 1 | 1 | 1 | PB1-111 | 1 | 1 | 1 | PA-641 | 1 | 1 | 1 | HA-563 | 1 | 1 | 1 | NP-107 | 1 | 1 | 1 | NA-687 | 1 | 1 | 1 | M-351 | 1 | 1 | 1 | NS-530 | 1 | 1 |
| **Minor77** | 10 | 1 | 1 | PB2-138 | 1 | 1 | 1 | PB1-673 | 1 | 1 | 1 | PA-606 | 1 | 1 | 1 | HA-562 | 1 | 1 | 1 | NP-467 | 1 | 1 | 1 | NA-681 | 1 | 1 | 1 | M-351 | 1 | 1 | 1 | NS-530 | 1 | 1 |
| **Minor98** | 10 | 2 | 1 | PB2-20 | 1 | 1 | 1 | PB1-631 | 1 | 1 | 1 | PA-40 | 1 | 1 | 1 | HA-445 | 1 | 1 | 1 | NP-100 | 1 | 1 | 1 | NA-172 | 1 | 1 | 1 | M-332 | 1 | 1 | 2 | NS-589–NS-592 | 4 | 2 |

**Table S6.** Phylogenetic cluster composition of Genin2 genotypes with at least 10 sequences. Clusters were numbered from bottom to top of trees. Number of sequences and number of TreeCluster (TC) genotypes; for each segment, the number of TreeCluster clusters, range of cluster IDs, span of cluster IDs (max-min+1), and the ratio of span to number of clusters. Number and range of clusters exclude singletons; number of genotypes (genos) includes singletons.

|  |  |  | **PB2** | | | | **PB1** | | | | **PA** | | | | **HA** | | | | **NP** | | | | **NA** | | | | **M** | | | | **NS** | | | |
| --- | --- | --- | --- | --- | --- | --- | --- | --- | --- | --- | --- | --- | --- | --- | --- | --- | --- | --- | --- | --- | --- | --- | --- | --- | --- | --- | --- | --- | --- | --- | --- | --- | --- | --- |
| **Genin**  **genotype** | N seq | N TC geno | N clust | ID range | Span | Span  /N | N clust | ID range | Span | Span  /N | N clust | ID range | Span | Span  /N | N clust | ID range | Span | Span  /N | N clust | ID range | Span | Span  /N | N clust | ID range | Span | Span  /N | N clust | ID range | Span | Span  /N | N clust | ID range | Span | Span  /N |
| **EA-2020-C** | 3361 | 240 | 7 | PB2-180 – PB2-186 | 7 | 1 | 12 | PB1-153 – PB1-673 | 521 | 43.4 | 30 | PA-587 – PA-631 | 45 | 1.5 | - | - | - | - | 19 | NP-466 – NP-484 | 19 | 1 | 27 | NA-638 – NA-704 | 67 | 2.5 | 32 | M-316 – M-361 | 46 | 1.4 | 29 | NS-530 – NS-592 | 63 | 2.2 |
| **EA-2022-BB** | 1758 | 60 | 12 | PB2-661 – PB2-672 | 12 | 1 | 18 | PB1-655 – PB1-672 | 18 | 1 | 2 | PA-461 – PA-462 | 2 | 1 | - | - | - | - | 11 | NP-117 – NP-127 | 11 | 1 | 1 | NA-675 | 1 | 1 | 1 | M-337 | 1 | 1 | 3 | NS-461 – NS-463 | 3 | 1 |
| **EA-2020-A** | 1751 | 218 | 6 | PB2-604 – PB2-609 | 6 | 1 | 19 | PB1-549 – PB1-597 | 49 | 2.6 | 16 | PA-538 – PA-553 | 16 | 1 | - | - | - | - | 21 | NP-422 – NP-442 | 21 | 1 | 5 | NA-281 – NA-289 | 9 | 1.8 | 40 | M-263 – M-315 | 53 | 1.3 | 5 | NS-394 – NS-398 | 5 | 1 |
| **EA-2024-DI** | 1528 | 5 | 1 | PB2-645 | 1 | 1 | 1 | PB1-627 | 1 | 1 | 4 | PA-581 – PA-584 | 4 | 1 | - | - | - | - | 1 | NP-463 | 1 | 1 | 1 | NA-671 | 1 | 1 | 1 | M-338 | 1 | 1 | 1 | NS-453 | 1 | 1 |
| **EA-2021-AB** | 1430 | 155 | 8 | PB2-654 – PB2-671 | 18 | 2.2 | 17 | PB1-482 – PB1-654 | 173 | 10.2 | 8 | PA-565 – PA-573 | 9 | 1.1 | - | - | - | - | 8 | NP-456 – NP-463 | 8 | 1 | 6 | NA-670 – NA-675 | 6 | 1 | 7 | M-330 – M-338 | 9 | 1.3 | 45 | NS-535 – NS-592 | 58 | 1.3 |
| **EA-2021-I** | 338 | 4 | 1 | PB2-575 | 1 | 1 | 1 | PB1-626 | 1 | 1 | 1 | PA-506 | 1 | 1 | - | - | - | - | 1 | NP-451 | 1 | 1 | 1 | NA-276 | 1 | 1 | 4 | M-284 – M-288 | 5 | 1.2 | 1 | NS-420 | 1 | 1 |
| **EA-2022-CH** | 150 | 9 | 4 | PB2-651 – PB2-659 | 9 | 2.2 | 2 | PB1-640 – PB1-653 | 14 | 7 | 6 | PA-566 – PA-573 | 8 | 1.3 | - | - | - | - | 2 | NP-450 – NP-451 | 2 | 1 | 3 | NA-671 – NA-675 | 5 | 1.7 | 1 | M-338 | 1 | 1 | 3 | NS-553 – NS-589 | 37 | 12.3 |
| **EA-2023-DA** | 121 | 6 | 1 | PB2-576 | 1 | 1 | 1 | PB1-485 | 1 | 1 | 4 | PA-574 – PA-580 | 7 | 1.8 | - | - | - | - | 1 | NP-463 | 1 | 1 | 1 | NA-671 | 1 | 1 | 1 | M-338 | 1 | 1 | 2 | NS-452 – NS-453 | 2 | 1 |
| **EA-2021-AF** | 47 | 5 | 1 | PB2-216 | 1 | 1 | 1 | PB1-485 | 1 | 1 | 2 | PA-591 – PA-596 | 6 | 3 | - | - | - | - | 1 | NP-448 | 1 | 1 | 1 | NA-663 | 1 | 1 | 4 | M-324 – M-330 | 7 | 1.8 | 2 | NS-20 – NS-21 | 2 | 1 |
| **EA-2023-DT** | 46 | 1 | 1 | PB2-672 | 1 | 1 | 1 | PB1-154 | 1 | 1 | 1 | PA-462 | 1 | 1 | - | - | - | - | 1 | NP-127 | 1 | 1 | 1 | NA-675 | 1 | 1 | 1 | M-337 | 1 | 1 | 1 | NS-463 | 1 | 1 |
| **EA-2023-DG** | 44 | 3 | 1 | PB2-553 | 1 | 1 | 1 | PB1-653 | 1 | 1 | 1 | PA-561 | 1 | 1 | - | - | - | - | 1 | NP-460 | 1 | 1 | 1 | NA-674 | 1 | 1 | 1 | M-333 | 1 | 1 | 3 | NS-586 – NS-588 | 3 | 1 |
| **EA-2023-DB** | 37 | 5 | 1 | PB2-658 | 1 | 1 | 3 | PB1-482 – PB1-653 | 172 | 57.3 | 1 | PA-566 | 1 | 1 | - | - | - | - | 2 | NP-460 – NP-462 | 3 | 1.5 | 2 | NA-674 – NA-675 | 2 | 1 | 2 | M-333 – M-334 | 2 | 1 | 4 | NS-550 – NS-588 | 39 | 9.8 |
| **EA-2021-AD** | 32 | 1 | 1 | PB2-577 | 1 | 1 | 1 | PB1-601 | 1 | 1 | 1 | PA-587 | 1 | 1 | - | - | - | - | 1 | NP-452 | 1 | 1 | 1 | NA-668 | 1 | 1 | 1 | M-320 | 1 | 1 | 1 | NS-592 | 1 | 1 |
| **EA-2021-O** | 32 | 1 | 1 | PB2-609 | 1 | 1 | 1 | PB1-504 | 1 | 1 | 1 | PA-561 | 1 | 1 | - | - | - | - | 1 | NP-433 | 1 | 1 | 1 | NA-285 | 1 | 1 | 1 | M-309 | 1 | 1 | 1 | NS-398 | 1 | 1 |
| **EA-2022-CC** | 27 | 6 | 1 | PB2-216 | 1 | 1 | 1 | PB1-653 | 1 | 1 | 1 | PA-503 | 1 | 1 | - | - | - | - | 1 | NP-462 | 1 | 1 | 1 | NA-675 | 1 | 1 | 1 | M-338 | 1 | 1 | 5 | NS-542 – NS-549 | 8 | 1.6 |
| **EA-2024-EF** | 19 | 1 | 1 | PB2-645 | 1 | 1 | 1 | PB1-627 | 1 | 1 | 1 | PA-581 | 1 | 1 | - | - | - | - | 1 | NP-449 | 1 | 1 | 1 | NA-671 | 1 | 1 | 1 | M-338 | 1 | 1 | 1 | NS-453 | 1 | 1 |
| **EA-2020-E** | 17 | 8 | 1 | PB2-609 | 1 | 1 | 1 | PB1-563 | 1 | 1 | 1 | PA-560 | 1 | 1 | - | - | - | - | 1 | NP-442 | 1 | 1 | 3 | NA-266 – NA-268 | 3 | 1 | 5 | M-278 – M-313 | 36 | 7.2 | 1 | NS-398 | 1 | 1 |
| **EA-2020-G** | 17 | 6 | 1 | PB2-646 | 1 | 1 | 1 | PB1-617 | 1 | 1 | 1 | PA-557 | 1 | 1 | - | - | - | - | 1 | NP-350 | 1 | 1 | 1 | NA-18 | 1 | 1 | 6 | M-281 – M-313 | 33 | 5.5 | 1 | NS-529 | 1 | 1 |
| **EA-2021-Q** | 17 | 7 | 2 | PB2-596 – PB2-609 | 14 | 7 | 2 | PB1-536 – PB1-565 | 30 | 15 | 2 | PA-526 – PA-553 | 28 | 14 | - | - | - | - | 2 | NP-417 – NP-451 | 35 | 17.5 | 2 | NA-285 – NA-294 | 10 | 5 | 4 | M-264 – M-313 | 50 | 12.5 | 4 | NS-378 – NS-398 | 21 | 5.2 |
| **EA-2021-BA** | 15 | 5 | 4 | PB2-171 – PB2-178 | 8 | 2 | 2 | PB1-632 – PB1-673 | 42 | 21 | 2 | PA-596 – PA-603 | 8 | 4 | - | - | - | - | 3 | NP-467 – NP-474 | 8 | 2.7 | 3 | NA-663 – NA-680 | 18 | 6 | 3 | M-330 – M-346 | 17 | 5.7 | 4 | NS-531 – NS-589 | 59 | 14.8 |
| **EA-2022-AM** | 12 | 3 | 1 | PB2-186 | 1 | 1 | 1 | PB1-632 | 1 | 1 | 1 | PA-596 | 1 | 1 | - | - | - | - | 1 | NP-350 | 1 | 1 | 1 | NA-663 | 1 | 1 | 2 | M-324 – M-330 | 7 | 3.5 | 2 | NS-557 – NS-567 | 11 | 5.5 |
| **EA-2023-DJ** | 12 | 1 | 1 | PB2-653 | 1 | 1 | 1 | PB1-485 | 1 | 1 | 1 | PA-576 | 1 | 1 | - | - | - | - | 1 | NP-451 | 1 | 1 | 1 | NA-671 | 1 | 1 | 1 | M-338 | 1 | 1 | 1 | NS-409 | 1 | 1 |
| **EA-2021-R** | 11 | 1 | 1 | PB2-609 | 1 | 1 | 1 | PB1-563 | 1 | 1 | 1 | PA-560 | 1 | 1 | - | - | - | - | 1 | NP-448 | 1 | 1 | 1 | NA-267 | 1 | 1 | 1 | M-314 | 1 | 1 | 1 | NS-398 | 1 | 1 |
| **EA-2024-EE** | 11 | 1 | 1 | PB2-645 | 1 | 1 | 1 | PB1-627 | 1 | 1 | 1 | PA-583 | 1 | 1 | - | - | - | - | 1 | NP-451 | 1 | 1 | 1 | NA-671 | 1 | 1 | 1 | M-338 | 1 | 1 | 1 | NS-453 | 1 | 1 |
| **EA-2021-AH** | 10 | 1 | 1 | PB2-216 | 1 | 1 | 1 | PB1-673 | 1 | 1 | 1 | PA-603 | 1 | 1 | - | - | - | - | 1 | NP-467 | 1 | 1 | 1 | NA-680 | 1 | 1 | 1 | M-346 | 1 | 1 | 1 | NS-531 | 1 | 1 |

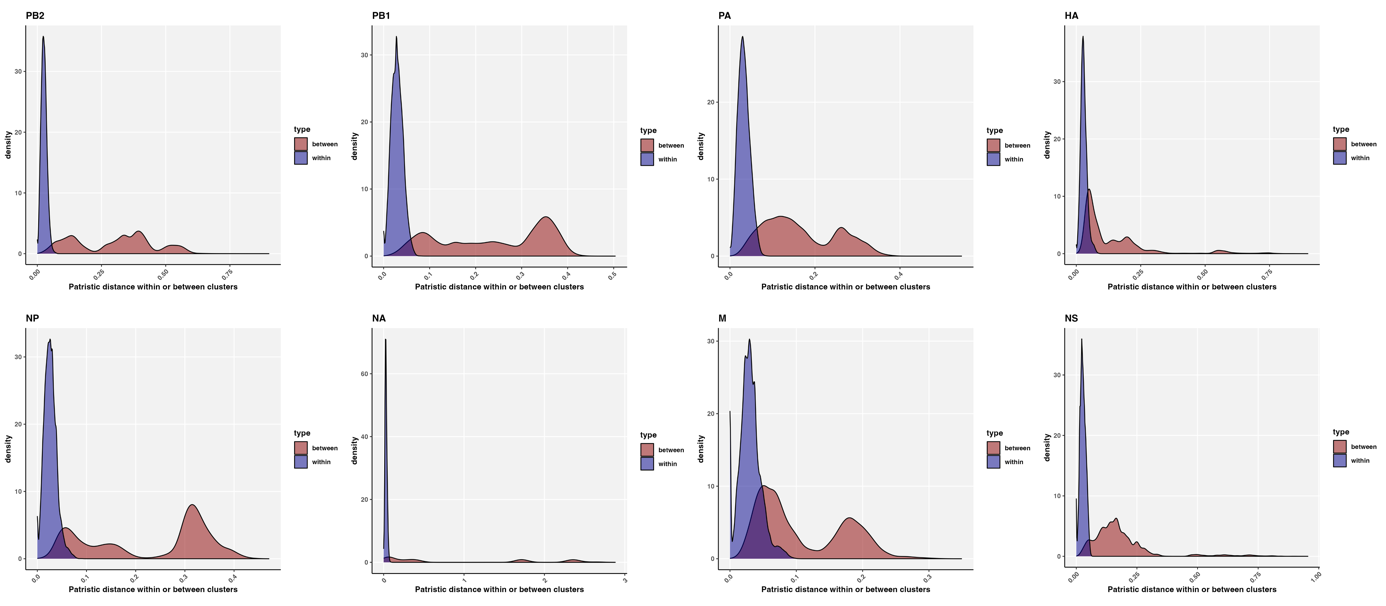

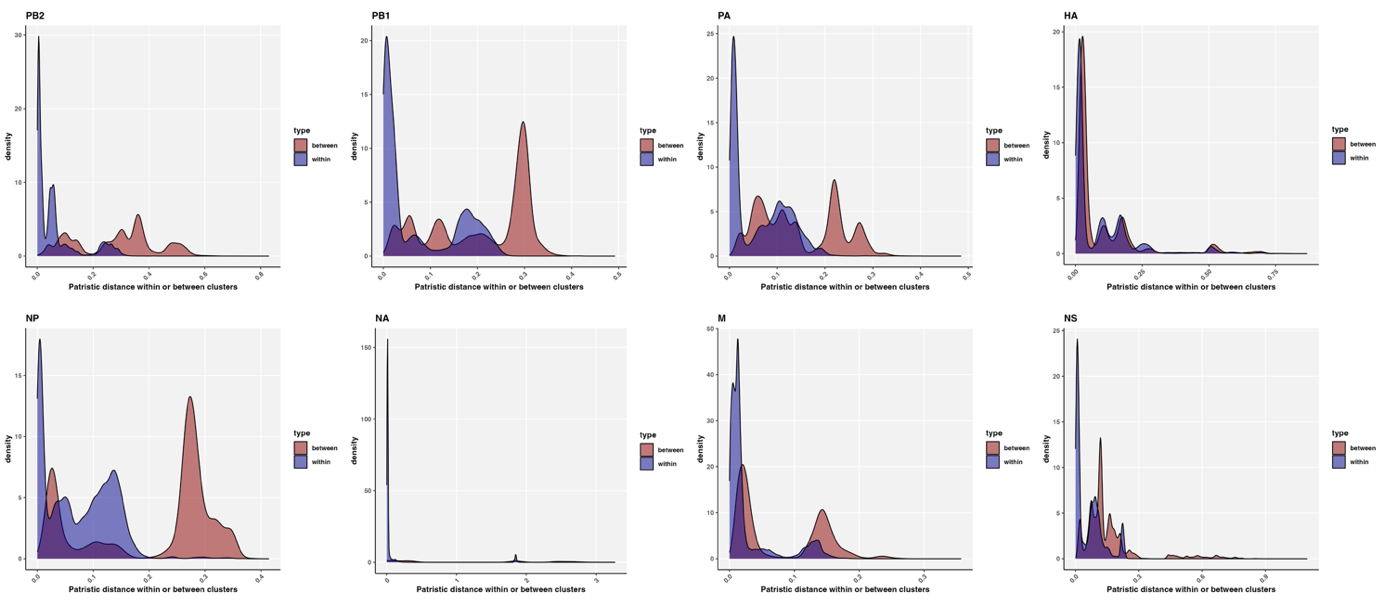
**Fig. S15.** Distributions of pairwise patristic distances (subs/site) within or between GenoFlu clusters.

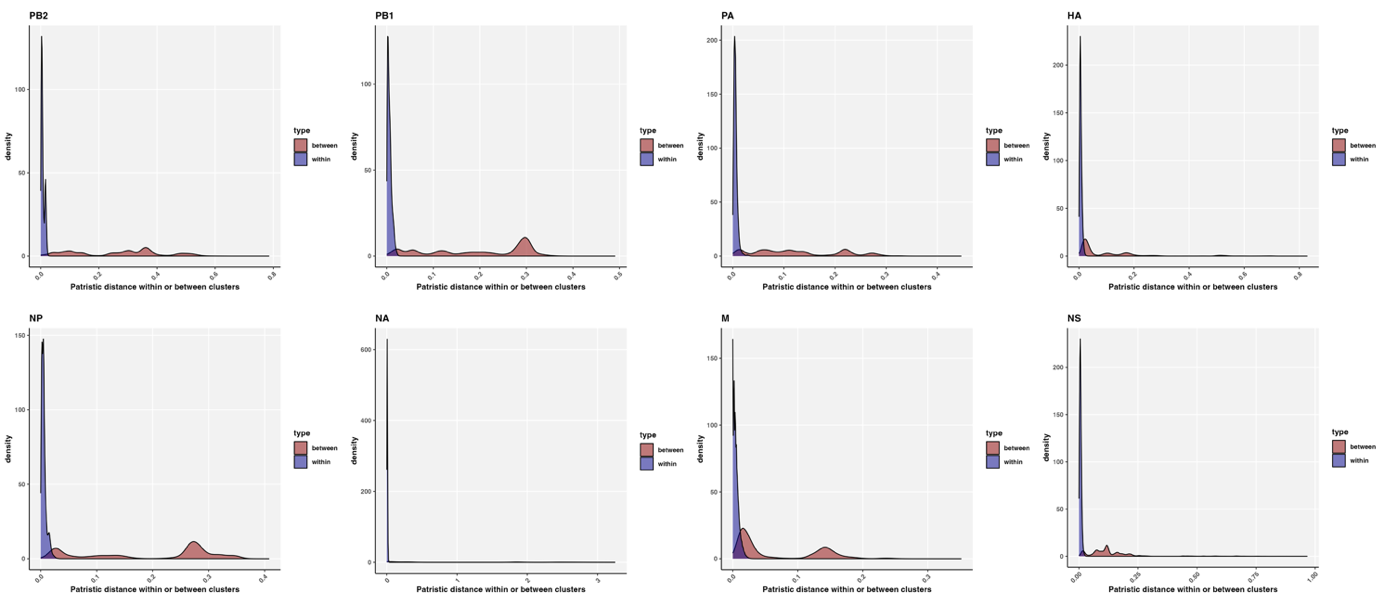
**Fig. S16.** Distributions of pairwise patristic distances within or between CD-HIT clusters (98%).

**Fig. S17.** Distributions of pairwise patristic distances within or between TreeClusters (0.04 subs/site).

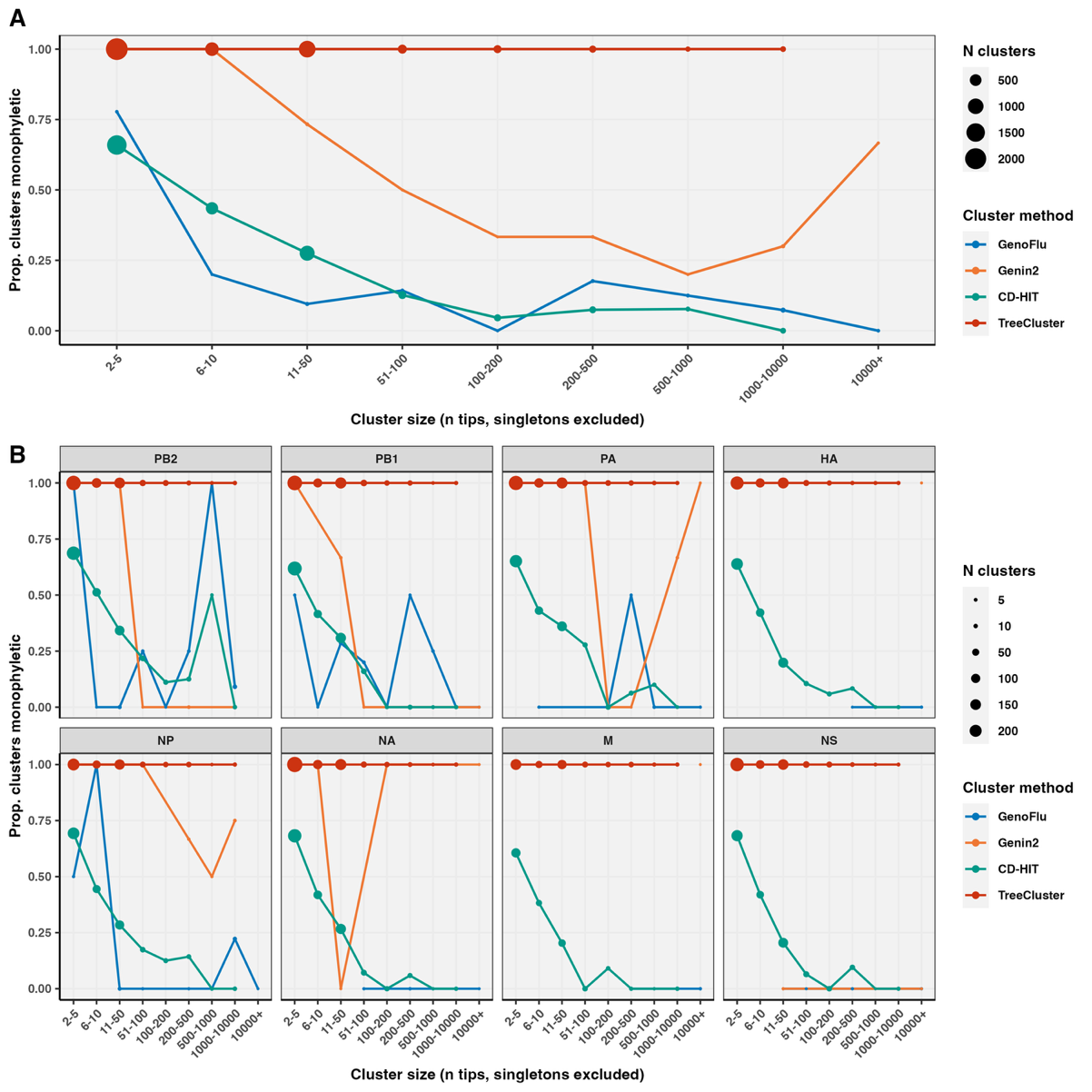
**Fig. S18.** Proportion of monophyletic clusters across cluster size, method, and segment. A) Overall proportion of monophyletic clusters of all segments, and B) by segment. Point size proportional to the number of clusters in size categories. Singletons and N/A labels excluded.

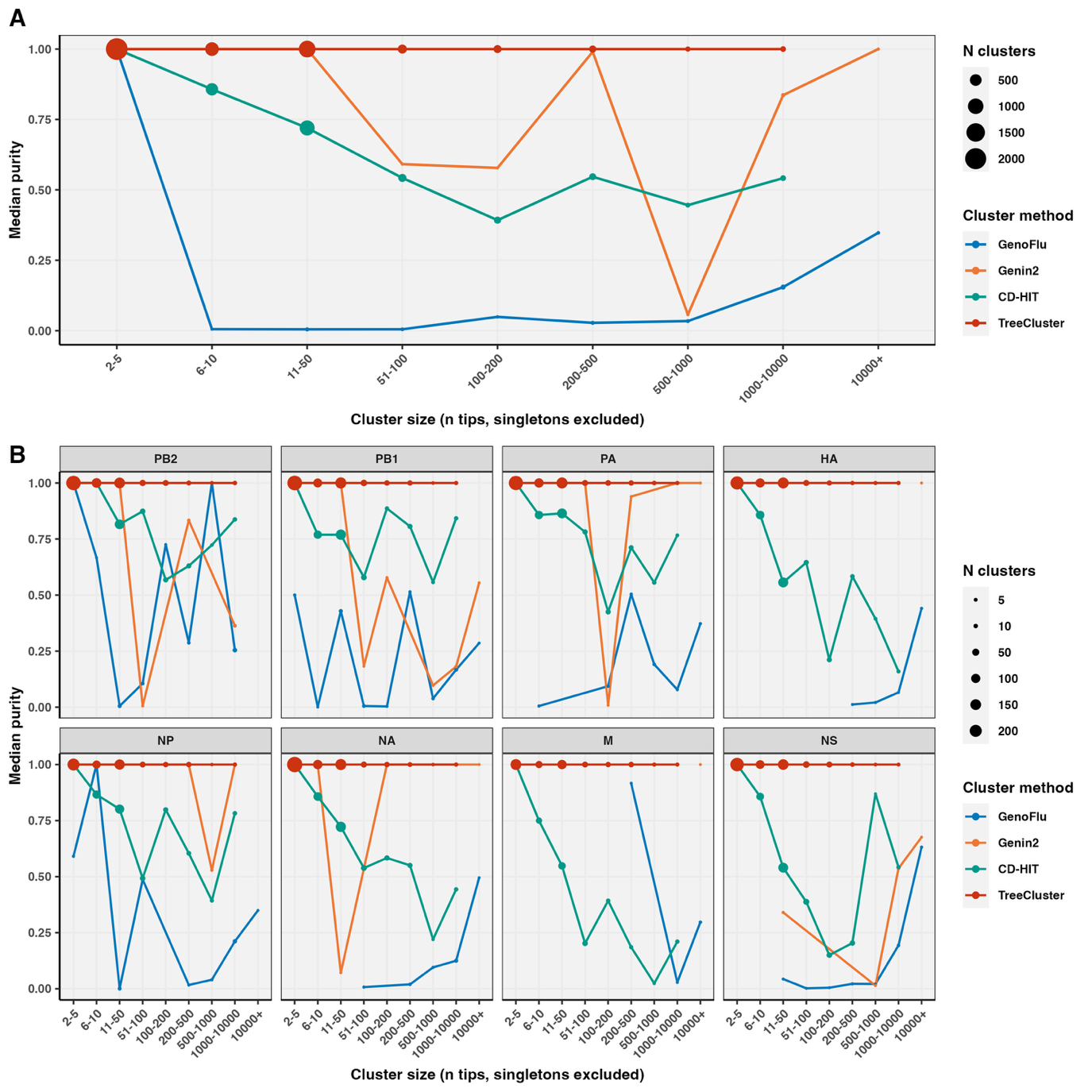
**Fig. S19.** Clusters’ median purity across cluster size, method, and segment. A) Median purity for clusters overall for all segments, and B) by segment. Point size proportional to the number of clusters in size categories. Singletons and N/A labels excluded.

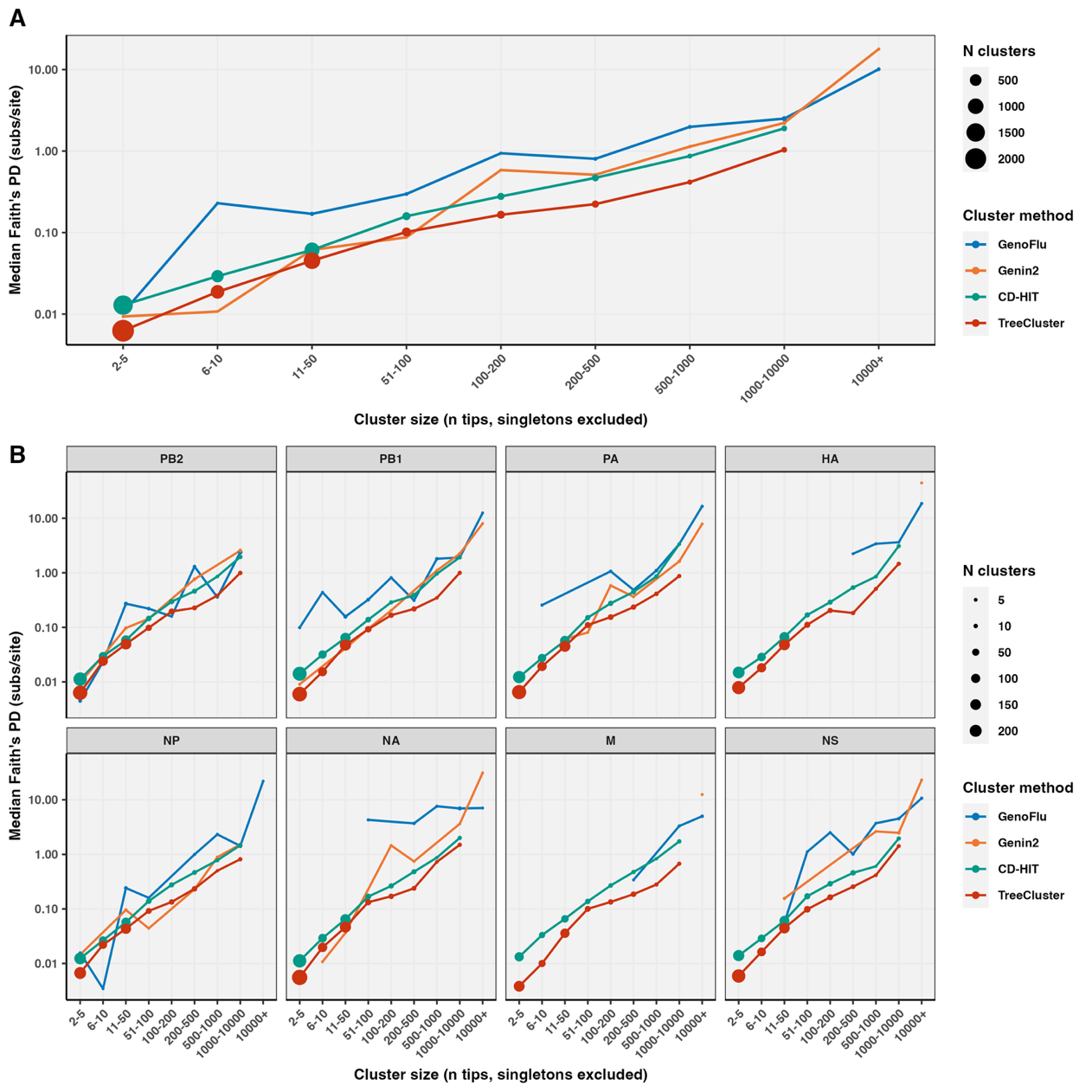
**Fig. S20.** Clusters’ median Faith’s phylogenetic diversity (PD) across cluster size, method, and segment. A) Median PD for clusters overall for all segments, and B) by segment. Point size proportional to the number of clusters in size categories. Singletons and N/A labels excluded.

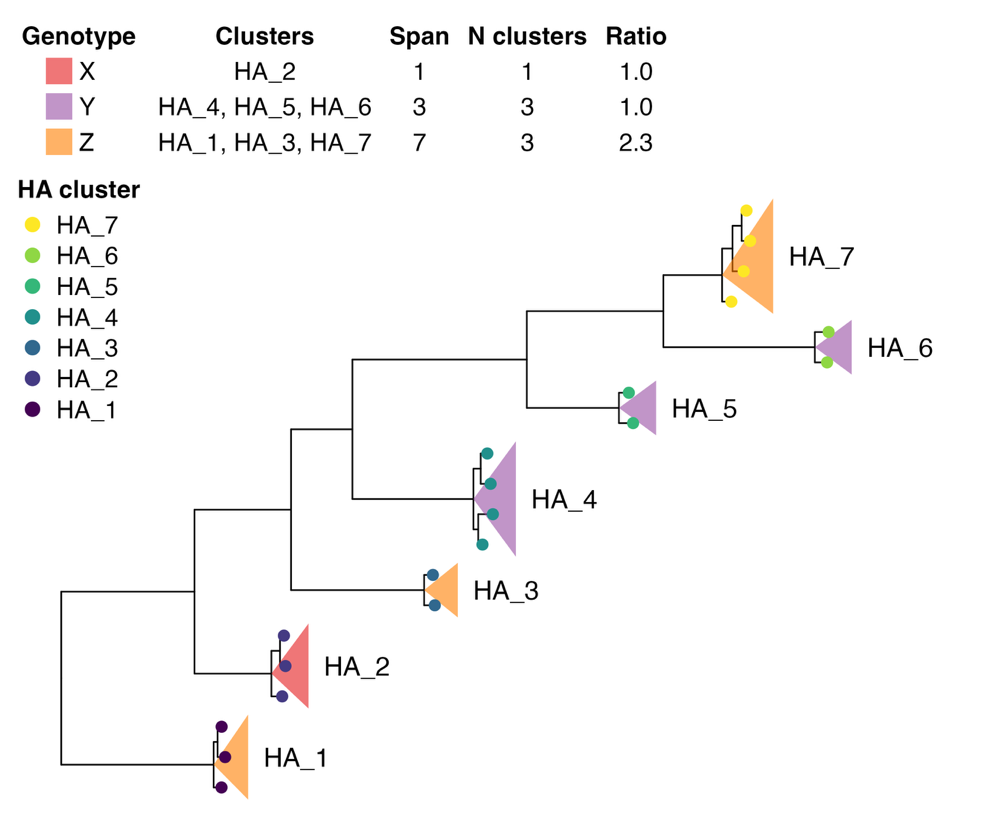

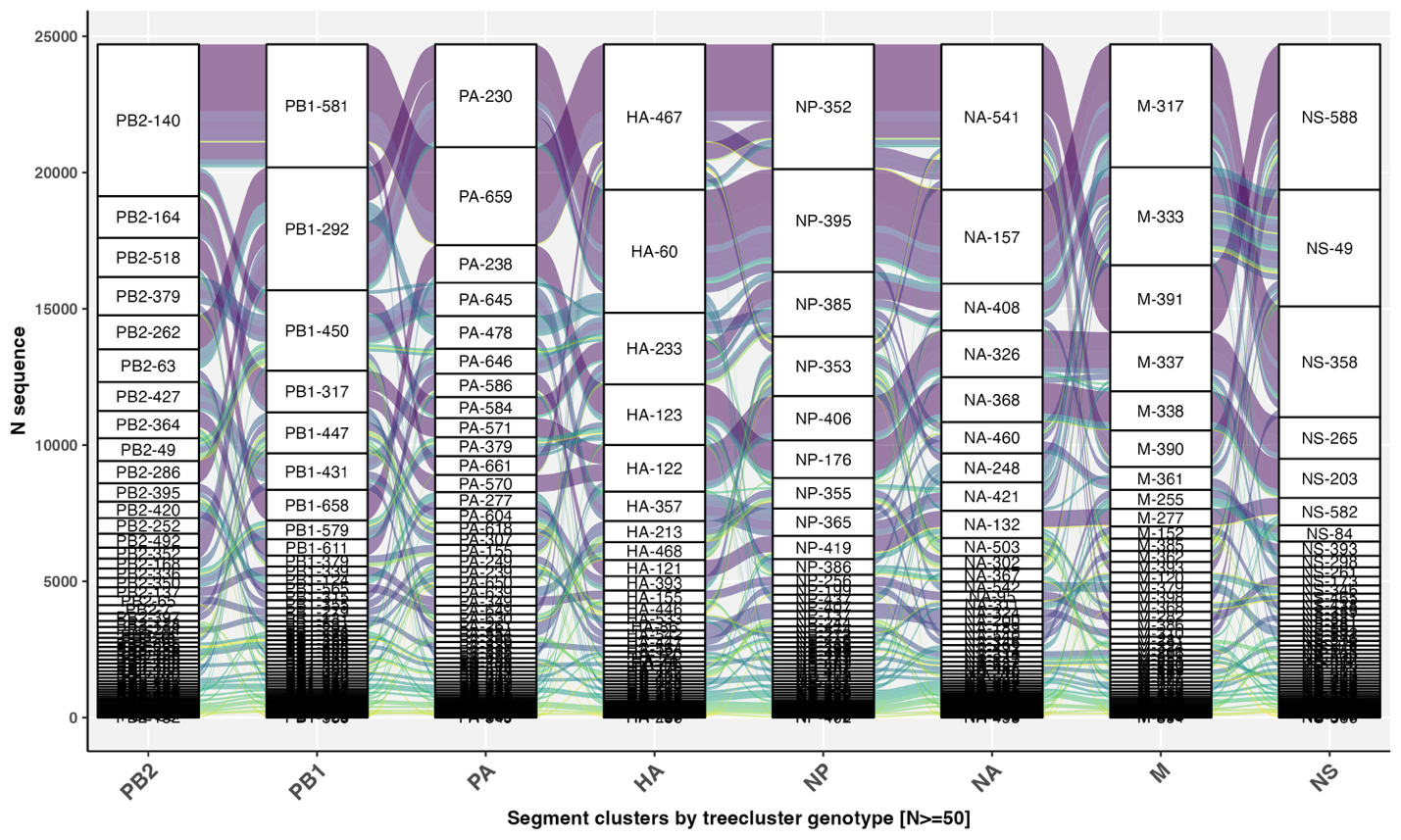
**Fig. S21**. H5Nx TreeCluster genotypes connect unique combinations of segment clusters. Only genotypes with at least 50 sequences shown.

**Fig. S22**. Phylogenetic cluster span ratio within predominant genotypes to distinguish drift from reassortment. Cluster span = maximum cluster ID – minimum cluster ID + 1; Span-to-cluster ratio = cluster span / number of clusters; where a ratio of 1 suggests clusters diversified by drift because they are contiguous, while a ratio exceeding 1 suggests the genotype may include multiple reassortants.
