## Supplementary material for "Influenza A virus H5Nx genotypes assigned by genetic similarity are incongruent with global phylogenies": GISAID supplementary file

### Supplementary Appendix

All genome sequences and associated metadata supporting the findings of this study can be accessed through the persistent digital object identifier https://doi.org/10.55876/gis8.260622ts

In addition to the minted DOI, GISAID also communicates the aggregation of GISAID accession numbers (EPI_ISL_IDs) through the corresponding EPI_SET_260622ts identifier to facilitate both, the acknowledgment of all data contributors and the direct retrieval of the underlying data from GISAID used in this study.

#### Influenza Virus Data Summary

| GISAID Identifier | Digital Object Identifier | Number of individual viruses | Data Collection range | Number of countries/territories |
|---|---|---|---|---|
| EPI_SET_260622ts | https://doi.org/10.55876/gis8.260622ts | 30,735 | 1975-01-01 to 2025-11-30 | 114 |
